# Seeing Above and Below the Canopy: Modeling and Interpreting Species Occupancy with Multimodal Habitat Representations

**DOI:** 10.1101/2025.09.06.674602

**Authors:** Timm Haucke, Lauren Harrell, Yunyi Shen, Levente Klein, David Rolnick, Lauren E. Gillespie, Sara Beery

## Abstract

Effective conservation and restoration of species is an increasingly urgent priority. To design management strategies that improve species success, we need a solid understanding of the habitat characteristics that support it. Occupancy models are statistical tools that ecologists use to model these relationships from data. Yet, current models represent habitats with coarse-scale environmental variables that fail to capture important microhabitat features. We show that these limitations can be addressed by incorporating AI-derived, multimodal habitat representations from overhead satellite imagery and ground-level camera-trap imagery. Across geography and species, these representations yield more accurate out-of-sample predictions than models based on conventional covariates alone, and combining satellite and ground-level views provides complementary gains. To translate improved prediction into actionable ecological insight, we further introduce a method that makes black-box AI-derived habitat representations interpretable by summarizing key factors contributing to occupancy probability into text-based descriptions. We then generate a per-site score for each description, which can replace black-box features to transparently link discovered habitat elements to species occurrence while maintaining predictive performance. Our approach provides a path toward microhabitat-aware and interpretable species-habitat models that support restoration planning and management decisions. We implement our method in an open-source Python package bridging AI and statistical ecology.

## 1 Introduction

Understanding what conditions drive where species occur is critical to understanding species niches [17], effective conservation planning [26], and human-wildlife conflict prevention [18]. Species distribution models (SDMs) use statistical approaches to link environmental variables such as temperature and elevation with observations of species’ presence, based on the hypothesis that a species’ occurrence is driven by the environmental characteristics of a region. However, observations of species presence do not paint the entire picture–a species may still be present even if unobserved.

Species occupancy models, a subset of SDMs, model this directly by using multiple observed detections and non-detections over time to explicitly account for imperfect detection [37]. By disentangling the probability of a species’ occupancy from the probability of its detection, species occupancy models produce less-biased estimates of where species occur [25]. Occupancy models have successfully been deployed to predict the risk of wildlife road crossings [55], detect invasive species early [6], identify high-priority conservation areas [13], and support harvesting decision-making [21].

The environmental variables predominantly used in existing occupancy models, e.g., average surface temperature, are often of coarse spatial and temporal resolution [5, 20], which can limit their predictive power [35]. Recent advances in SDMs have sought to address this limitation by using deep learning models to directly predict species distributions and encounter rates from higher-resolution satellite imagery [61, 47, 23, 56, 32].

While promising, the use of satellite imagery alone still fails to capture important aspects of the microhabitat, such as structure and composition of vegetation under the tree canopy of forests [24]. These microhabitat features *can* be captured in images taken on the ground [58]. Among the most widely deployed sources of such ground-level imagery are camera traps, motion-triggered cameras mounted on trees or posts for long periods of time that continuously and non-invasively detect species, which can provide unbiased [11] and systematic [53] species observations that are well-suited for species occupancy modeling. These ground-level images *also* capture microhabitat features challenging to see from space, like plant phenology timing [30] and leaf litter [60].

In this work, we propose the first scalable mechanism to enable these complementary but distinct habitat features (from environmental variables, satellites, and ground-level images) to inform our models of species occupancy. High-level internal representations of raw images generated by deep learning models provide a way to extract microhabitat clues. Importantly, this does not require prior knowledge about which cues are relevant, nor does it require labeling such cues manually. By utilizing the resulting multimodal deep features as covariates within a Bayesian occupancy modeling method, we obtain models that make use of fine-grained habitat features while maintaining the statistical rigor of detection/non-detection inference.

We demonstrate the value of multimodal deep features by fitting occupancy models on large-scale data from the Wildlife Insights platform [3]. Evaluating the fitted models on publicly-available camera trap data from the nationwide Snapshot USA camera trapping surveys [53] reveals that these multimodal deep features significantly improve occupancy modeling accuracy for a clear majority of modeled species.

A challenge of using deep features as predictors is that they are difficult to interpret. Whereas traditional ecological models predominantly learn few coefficients that directly correspond to environmental variables, our deep multimodal models contain hundreds of coefficients that correspond to a priori uninterpretable features. To overcome the inherent black-box nature of deep features in the context of occupancy modeling, we propose an interpretability technique that summarizes in natural language habitat elements that the models deem important. These habitat elements can then be compared to prior knowledge and converted into simplified habitat covariates that condense information contained within images, thereby regaining the ability to interpret model coefficients directly.

## 2 Results

### 2.1 Deep multimodal occupancy models

Our central question is whether imagery (e.g., from satellites or from camera traps) can act as additional useful environmental context for site-level occupancy models: can a model that already uses standard environmental covariates *X*_env_ gain predictive power when we augment those covariates with (i) satellite-derived embeddings *X*_sat_ and/or (ii) embeddings from a single blank (no-animal) daytime camera-trap image *X*_img_? To answer this question, we fit a suite of logit-linear occupancy models [37] (see Figure 1 and Methods 4.2 for occupancy model details) that differ only in which of these modalities they include (all combinations of *X*_env_, *X*_sat_, *X*_img_, see Methods 4.1). We then evaluate out-of-sample predictive performance on the detection histories of held-out sites using normalized log point-wise predictive density (LPPD, see Equation (4) and Methods 4.3).

**Figure 1:**
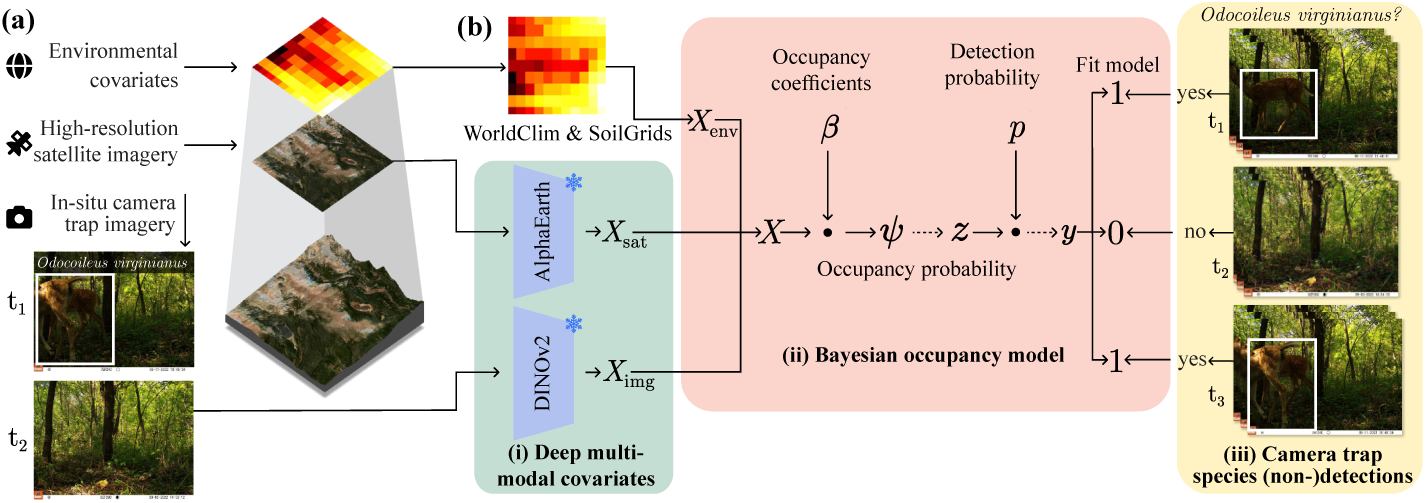
A deep multimodal species occupancy modeling method. **(a)** Ground-level camera trap blanks capture important microhabitat information like under-story structure [24], while high-resolution satellite imagery describes complementary site-level features not well-captured by local-scale traditional environmental covariates [45]. **(b)** Our deep multimodal species occupancy model is composed of: (i) deep microhabitat features derived from satellite imagery and camera trap blanks using earth observation and vision foundation models; (ii) Bayesian hierarchical occupancy models [37] implemented in a new deep learning-optimized Python package called *Biolith* (see section C.2); and (iii) species detection and non-detection data derived from the same camera trap imagery used to derive microhabitat features. Example camera trap images are from Snapshot USA 2022 [53].

To make LPPD easier to interpret and compare across species, we normalize each species’ test-set LPPD to a scale defined by a null (intercept-only) baseline and an “oracle” upper bound overfit to the test data (LPPD_norm_, see Equation (5)).

We train continental-scale occupancy models using 83 individual camera trap datasets sourced from Wildlife Insights and evaluate on a held-out Snapshot USA [53] test set (see Figure 4, Methods 4.1.1). To prevent data leakage and to limit the impact of spatial autocorrelation on evaluation, we additionally enforce a 10 km buffer between train and test sites. We utilize the occupancy likelihood as the primary evaluation target as it directly measures how well a model explains repeated detection / non-detection histories under imperfect detection.

### 2.2 Adding imagery improves predictive performance

Surprisingly, we see that simply scaling traditional environmental covariate-based occupancy modeling approaches to continental-scale data does not significantly improve performance over the null assumption of constant occupancy and detection rates estimated from Wildlife Insights data (non-significant *p*-value, one-sample Wilcoxon signed-rank test, one-tailed *H*_1_: LPPD_norm_ *>* 0). In contrast, any and all combinations of deep features consistently and significantly improve occupancy modeling performance over the null model (*p*-values 0.0467 −0.000381, one-tailed). This suggests that deep features are both sufficiently fine-grained yet generalized enough to describe how diverse species occupy similarly diverse habitats at continental scales.

Including satellite-and camera trap-derived deep features *together* significantly improves occupancy modeling performance over environmental covariates alone (paired Wilcoxon signed-rank test, one-tailed *H*_1_: 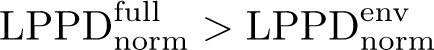 , Holm-adjusted *p* = 0.00092). The positive influence of multimodal deep features is additive, as including satellite imagery increases LPPD_norm_ by on average +0.10 (95% CI: [0.01 − 0.20], Tukey HSD-adjusted *p* = 0.031, see Methods 4.3.1) while including ground-level imagery increases LPPD_norm_ by on average +0.17 (95% CI: [0.07 − 0.26], Tukey HSD-adjusted *p* = 0.00054, 3-way ANOVA with Tukey’s Honest Significant Difference). Interaction effects were non-significant, suggesting that satellite and camera trap-derived features are each capturing complementary sources of the habitat information that drives species’ distributions. While both satellite and particularly ground-level imagery improve predictive performance on average, there is substantial species-to-species variation.

#### Fine-grained habitat structure: strong benefits from ground-level imagery

Several small mammals show large improvements when *X*_img_ is included, consistent with occupancy depending on microhabitat structure visible from the camera trap viewpoint. Predictive performance of the Eastern chipmunk (*Tamias striatus*), a small burrowing squirrel, improves from 0.18 with *X*_env_ to 0.40 with *X*_img_, and to 0.55 when *X*_sat_ and *X*_img_ are combined (Figure 2 and Supplemental Table B.1). The Eastern gray squirrel (*Sciurus carolinensis*) follows a similar pattern (0.20 → 0.49 with *X*_env_+*X*_img_, and 0.61 with *X*_sat_+*X*_img_). The Western gray squirrel (*Sciurus griseus*) shows the most benefit from ground-level imagery: environmental and satellite-only models fail to outperform the null model (−0.43 and −0.73), while ground-level imagery alone reaches 0.40. We hypothesize that this is because ground-level image encodes microhabitat details such as understory vegetation or deadwood, which are difficult to model using traditional environmental variables and difficult to assess by remote sensing, for example due to tree canopy occlusion.

**Figure 2:**
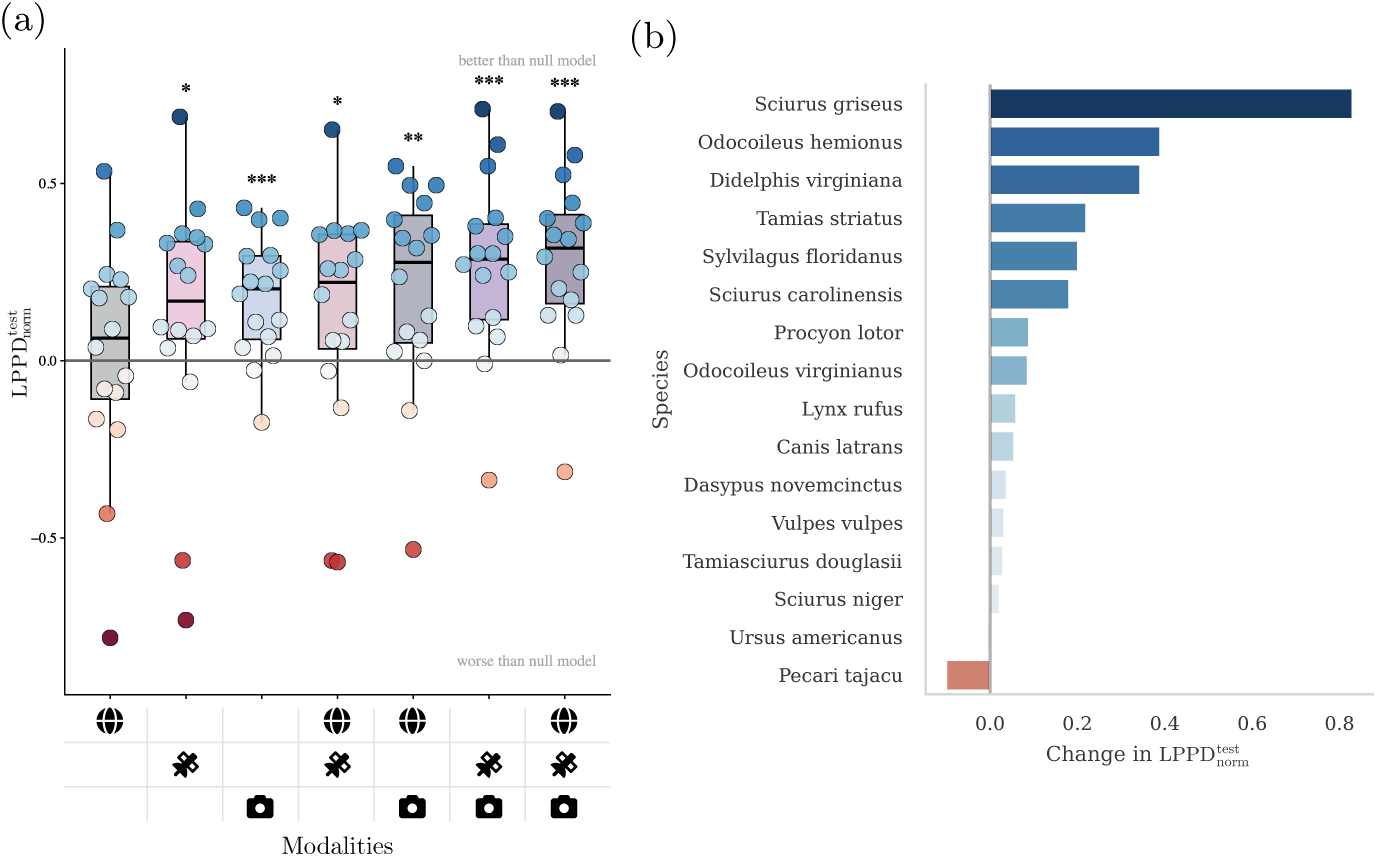
Satellite and ground-level imagery significantly improve predictive performance across a continental-scale occupancy modeling study. **(a)** Predictive performance summarized across species for different combinations of environmental variables 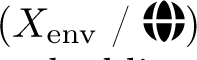, satellite imagery embeddings 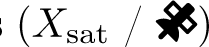, and ground-level imagery embeddings 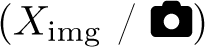. Points represent individual species. Stars indicate results from one-sample Wilcoxon signed-rank tests vs. the null model. *** indicates a *p*-value *<* 10^−3^, and * indicates a *p*-value *<* 10^−1^. **(b)** Change in predictive performance when adding ground-level imagery embeddings 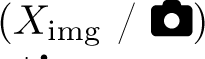. **Incorporating ground-level image features improves predictive power for the majority of species.** Improvement is measured for the best-performing model with ground-level image features over the best model without. Full results can be found in Supplemental Table B.1.

#### Generalists and carnivores: modality-specific results

For generalists and carnivores, imagery generally improves over environmental variables alone, but whether satellite or ground-level imagery helps more varies by species. The Northern raccoon (*Procyon lotor*) improves from 0.09 with *X*_env_ to 0.44 with the full multimodal model, and the combined *X*_sat_+*X*_img_ configuration is already near-optimal (Figure 2 and Supplemental Table B.1). The Bobcat (*Lynx rufus*) improves more modestly but most strongly when ground-level imagery is included (0.18 with *X*_env_ versus 0.24 with *X*_env_+*X*_img_), consistent with local cover and movement corridors being potentially informative [1]. The American black bear (*Ursus americanus*) provides a counterexample where satellite context appears to dominate: the best-performing configuration uses *X*_sat_ alone (0.36), and adding ground-level imagery does not improve performance. One possible interpretation is that American black bear occupancy in our datasets is more associated with broad landscape characteristics that satellite embeddings capture sufficiently well, while microhabitat specifics add comparatively little signal.

#### Ungulates: mixed results and distribution shift

White-tailed deer (*Odocoileus virginianus*) benefits from imagery, particularly when paired with environmental variables (0.44 for *X*_env_+*X*_img_), indicating that microhabitat context augments coarse predictors for this species. Mule deer (Odocoileus hemionus), however, represents an outlier: none of the candidate predictor sets outperform the null model (best score −0.17), which is likely caused by a significant distribution shift between our train and test splits.

### 2.3 Interpreting and simplifying complex models

Predictive improvements are most useful when they can be translated into ecological understanding, which is important for studying the ecology of species [17], decision making [57], and conservation [13]. In classic environmental-variable-based logit-linear occupancy models, regression coefficients corresponding to each environmental variable can be directly interpreted as estimating the direction and magnitude of influence on species occupancy or detection probability. In contrast, models using deep features as predictors are not initially interpretable, since each individual feature does not directly correspond to an a priori interpretable quantity. Since these deep features are in general difficult to interpret, deep models are frequently called “black-boxes”. Despite the black-box nature of deep representations, we can still build methods around the black-box to try and explain how the inputs influence its outputs. Hence, we propose a technique (Figure 3) that turns fitted multimodal occupancy models into (i) qualitative habitat elements, and (ii), a compact, interpretable set of covariates that can replace high-dimensional image embeddings. This technique can be broadly categorized into three phases.

**Figure 3:**
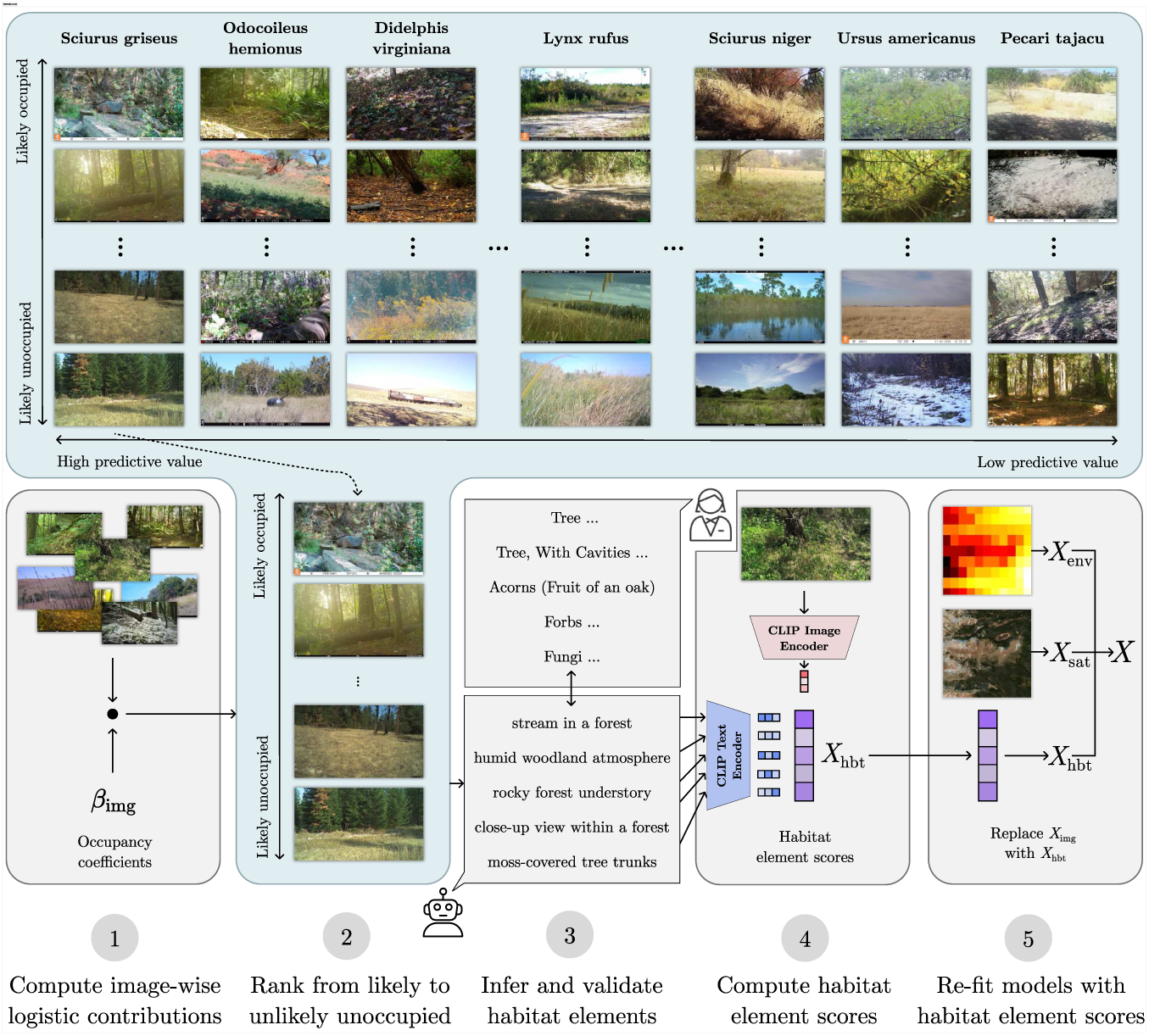
Method to automatically discover microhabitat elements from images. We start with an occupancy model that has learned some relationships between ground-level image features *X*_img_ and occupancy probability *ψ*, captured by the relevant model coefficients *β*_img_. We then rank each image from most likely occupied to most likely unoccupied, which provides a qualitative way to evaluate which microhabitat features the model is focusing on. Using an adaptation of VisDiff [16], we then summarize a set of five habitat elements in natural language that describe elements present in the likely occupied, but absent in the likely unoccupied images. These habitat elements can then be compared to prior knowledge, facilitating expert validation. Finally, we use CLIP [50] to quantify the presence of each habitat feature across all images, resulting in five-dimensional covariates. By replacing the high-dimensional ground-level image features with these five covariates described in natural language, we make our models interpretable without sacrificing predictive performance.

First, we rank ground-level images by their logistic contribution to occupancy using the dot product 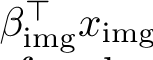. This yields intuitive image rankings for many species. For example, for the Western gray squirrel, the top-ranked images emphasize dense forest and deadwood (Figure 3).

Second, we adapt VisDiff [16] to summarize differences between likely-occupied and likely-unoccupied image sets as natural-language habitat elements (see Methods 4.4 for details). Species-specific examples illustrate these automatically extracted hypotheses. For Douglas’ squirrel (*Tamiasciurus douglasii*), extracted elements emphasize shaded coniferous forest microhabitat (e.g., mossy logs, fern-rich forest floor, dense understory), while expert-curated habitat elements emphasize mature conifer forest and associated structural cover (Supplemental Table B.4). For the bobcat, descriptions focus on elements that can act as movement corridors, such as dirt roads or open trails through forest and leaf-litter ground cover (Supplemental Table B.4). These descriptions are not intended to be exhaustive habitat models; rather, they provide image-grounded hypotheses that can and should be checked against expert knowledge and used to prioritize and generate targeted covariates.

In the third phase, we demonstrate (with additional creative use of multimodal AI models) that **these derived and discovered features can be strong predictors**. We use the CLIP [50] vision-language model (see Methods 4.4) to generate a continuous score between -1 and 1 capturing the presence of each of the top *k* = 5 derived natural language habitat elements across all ground-level images. This results in a compact, easily-interpretable *k*-dimensional continuous covariate *X*_hbt_. To evaluate the predictive power of *X*_hbt_, we re-fit our full multimodal occupancy models with these compact features in place of the ground-level image embeddings *X*img. We find that using *X*_hbt_ largely maintains or even improves predictive performance when compared with the original, much higher-dimensional ground-level image features *X*img (average drop of 0.05 test-set normalized LPPD across species, for details see section B.3). This indicates that the habitat covariates indeed capture meaningful microhabitat information, while reducing overfitting induced by spurious correlations between the full ground-level image features and occupancy. We draw analogies to covariate selection in traditional statistical ecology [19].

Finally, we find that our habitat covariates capture significant unique information that is not predictable from remote sensing features alone (average *R*^2^ across species-specific models is 0.01).

### 2.4 Robustness across representations and model formulations

For the main results presented above, we use AlphaEarth Foundations [7] for extracting features from satellite imagery, and DINOv2 [42] for ground-level imagery. Across all ground-level and satellite feature extractors evaluated in Section B.2, we find that these choices perform best, in general, on both the train and test data. Still, many alternative backbones improve over non-imagery baselines for a substantial fraction of species, and performance differences between leading representations are largely insignificant, as summarized in Supplemental Tables B.2 and B.3.

Second, our findings are consistent under a simplified “naive” occupancy framing. Many focal species have relatively high posterior mean detection probabilities (Supplemental Figure B.1), motivating a complementary experiment where sites are labeled occupied if the species is detected at least once. Under naive occupancy, the relative predictive power of the different modalities broadly match the hierarchical occupancy model results (details in Section B.1), suggesting that the observed improvements in predictive performance by using imagery are not an artifact of a particular model specification.

## 3 Discussion

Our proposed deep multimodal occupancy modeling approach offers several key advantages. First, our method can directly make use of complementary fine-grained habitat details derived from ground-level imagery without additional labeling. Second, the use of deep features enables us to learn complex, and potentially previously unknown, relationships between species occupancy and environmental variables directly from the data, without requiring prior knowledge about, or access to data for, candidate covariates. Third, we show that our method improves predictive performance for 15 out of 16 focal species.

Finally, we propose a method to recover the interpretability of deep learning based occupancy models via summarizing relevant ground-level habitat elements in natural language and utilizing these summaries to construct sets of text-driven continuous-valued habitat covariates, which we demonstrate can maintain the increased predictive capacity of our direct deep-feature approach.

### Predictive power vs. interpretability: overcoming the need to choose

Traditional statistical ecology strongly emphasizes simple models with few predictors. Simple models have various advantages; variance tends to be lower (bias-variance tradeoff), models are quicker to fit; and, maybe most importantly, we can directly interpret model parameters. In a simple model, these parameters directly describe the influence of habitat features on species occurrence, and these relationships are used to inform conservation actions such as ecosystem regeneration. However, constraining ourselves to simple, interpretable models often comes at the expense of predictive performance [59, 40]. Because interpretability is so important for action, this has often led to a preference for interpretable models despite their lack of predictive capacity. But this begs the question: is it valuable to explain a model that is fundamentally non-descriptive of reality?

Our deep multimodal occupancy models initially seem diametrically opposite to the concept of simple and minimal models. Although we show that they are significantly more predictive, how is that useful if their parameters conceal direct habitat-occurrence relationships? We have proposed a new method (as seen in Section 2.3) that seeks to close the gap between predictive power and the interpretable knowledge of species-habitat relationships. We show that we can simplify these complex models by automatically distilling high-dimensional features into a set of human-interpretable text-based concepts that capture the key ground-level habitat features informing occurrence for a species. We further demonstrate the expressivity and importance of these features by proposing a method to transmit text-based simple habitat descriptions into continuous-scored covariates, which we show can maintain the predictive performance of the full ground-level image features when incorporated into our models. This approach gives us *both* simple interpretability and high predictive performance, but we believe it holds promise beyond that. It provides a method that enables us to leverage *any* open-ended habitat descriptions in natural language, for example those provided by local and/or academic experts, as usable, quantitative covariates within an occupancy model.

We are excited about the potential of this approach, particularly given the demonstrated performance across the set of species we consider. However, it is important to emphasize how the reliance on existing, pre-trained vision-language models may lead to biases [2]. The quantification, and possible mitigation, of these biases with respect to natural world imagery and species distributions is a clear avenue for future work.

### Alternative modalities and ecological models

In this study, we focus exclusively on occupancy modeling camera trap data. However, there are other ecological models where one could investigate incorporating deep features in addition to or in lieu of traditional covariates. Examples include presence-only SDMs (see Section A.1), N-mixture models (e.g., [54]), distance sampling models (e.g., [31]), and resource selection functions (e.g., [38]).

There are also other ecological data modalities that capture useful fine-grained ecological context alongside species occurrence, such as passive acoustic monitoring [10] or environmental DNA [9]. One can easily imagine taking a similar approach and using deep learning to represent and enable access to the ecological context of these samples. Blank images, the choice we made in this initial work, are an intuitive way to define those features for camera traps, but the use of a single blank image per location is just a starting point. There is additional, valuable environmental context in the temporal changes to that environment over the time period of deployment. The key challenge when expanding our approach, to handle temporal variation or to include additional modalities, is to define a meaningful and informative mechanism to generate a feature embedding that is representative of the microhabitat, and useful for the modeling task of interest (see Section B.2). It is an interesting avenue of future work to explore how to *optimally* generate these representative features for each modality, and for each of the many possible ecological models beyond occupancy.

### Disentangling occupancy and detection in ground-level imagery

A fundamental strength of occupancy models lies in their explicit separation of two ecological processes: the probability that a species occupies a site (*ψ*) and the probability of detecting that species given it is present (*p*). Ground-level camera trap imagery, however, captures visual information potentially informative to both processes simultaneously—and the boundary between what constitutes an “occupancy” vs. “detection” covariate is not always clear-cut.

Consider trails, which feature prominently in camera trap imagery due to common deployment strategies [34]. Whether trail presence should be modeled as influencing occupancy or detection is arguably species-dependent. For wide-ranging carnivores like bobcats, trails function as movement corridors integral to habitat selection [15], suggesting they are genuine microhabitat components influencing occupancy. For territorial small mammals, trails may have minimal influence on site occupancy but substantially affect the probability of photographing an individual crossing the camera’s field of view.

The temporal dimension adds further complexity. Detection probability varies with time of day, season, and weather—factors encoded both in image metadata and in visual cues like lighting, snow cover, and vegetation phenology. Our approach uses a single blank image per site, collapsing this temporal variation into a static representation, yet the same site may appear dramatically different across seasons with corresponding shifts in both detectability and effective occupancy.

In this work, we incorporated ground-level image embeddings from a single point in time and solely as occupancy covariates, while estimating constant detection probability across sites. This decision prioritized methodological simplicity but conflates features relevant to both processes. An exciting avenue for future work is to disentangle which visual features are most predictive of occupancy versus detection, for example by modeling detection using image-level embeddings from each survey occasion while modeling occupancy using aggregated site-level representations. Such approaches could yield both improved performance and deeper ecological insight into how microhabitat structure shapes species presence versus observability.

### Practical implications

While ground-level data, such as images and other modalities (see Section 3) can provide valuable microhabitat context, such data is also expensive to collect from scratch and at scale. Fortunately, sensors are decreasing in cost [63, 8, 4], cultural norms around data sharing in ecology are shifting [49, 51], and citizen science platforms such as iNaturalist or Xeno-canto are decentralizing the burden of ground-level data collection [14, 12]. As a result, growing volumes of such data are openly available. Our work suggests an additional, and valuable, opportunity for citizen scientists–ecologists seeking to improve access to fine-grained ecological context in a given area could prioritize “useful” habitat data collection, i.e. via bioblitzes [43, 39] targeting ecological context as opposed to species occurrence. What remains to be seen is *which* data is most useful when seeking to fill gaps and improve our understanding? Clearly, citizen scientists cannot be deployed everywhere on Earth, but what we learn through this process about which data is “most valuable” to collect on the ground can inform data prioritization strategies in remote areas, such as the Amazon or remote Arctic.

## 4 Methods

### 4.1 Data

As a test bed, we focus our experiments on data from Wildlife Insights [3], an online platform that users upload, label and automatically classify species in large volumes of camera trap images. On Wildlife Insights, each camera trap image is associated with a single camera trap site (which might have an associated geographic location), and each site is in turn associated with a single project. Projects might represent individual camera trapping surveys or be part of a larger collection of surveys.

Since ecological data such as species detections/non-detection histories frequently has spatial and temporal structure, it is critical to account for such structure when evaluating models in order not to violate independence assumptions and report overly optimistic results [52]. This issue is made even more critical when using deep features within a model, since their high-dimensionality can more easily lead models to rely on nuisance variables that have no relation to the ecological processes driving species occupancy. As an example, camera trap hardware within a single survey is frequently identical, and model-specific metadata embedded within each image could theoretically be used by models to learn “shortcuts” between camera trap model and species occurrence within the survey’s geographic region.

#### 4.1.1 Train / test split

To ensure our evaluation is not overly optimistic and enable fair comparisons between models using different modalities, we extrapolate our Wildlife Insights-trained models to a held-out “test set” of data from the public Snapshot USA camera trapping surveys from 2020 to 2023, also accessed through the Wildlife Insights platform. To prevent data leakage due to overlapping or visually similar satellite imagery, we furthermore remove any sites from the train set that are closer than 10 km to any test site.

We restrict each species’ analysis to sites within states where it is known to occur. This serves two purposes: it steers the model toward learning fine-scale habitat associations (rather than coarse geography), and it avoids fitting occupancy in clearly implausible areas, which can lead to unstable estimation. Using states as the filtering unit also aligns with many practical management decisions made by state agencies. For example, the Eastern gray squirrel is common in the eastern and midwestern U.S. but largely absent from much of the West despite seemingly suitable habitat; including western sites would mostly teach the model broad biogeographic absence rather than microhabitat structure. This type of range-based restriction is common in SDMs [27, 41], and the resulting inference should be interpreted as applying to that range.

Since our datasets include a long tail of rarely observed species that are difficult to model, we focus on species that are observed at least once at 10% of sites or more across their range. This results in 16 species that are reasonably prevalent within their range, some of which encompass close to the entire contiguous United States and some of which are highly localized. Figure 4 shows the spatial density of camera trap sites and summarizes species prevalence across our focal species.

**Figure 4:**
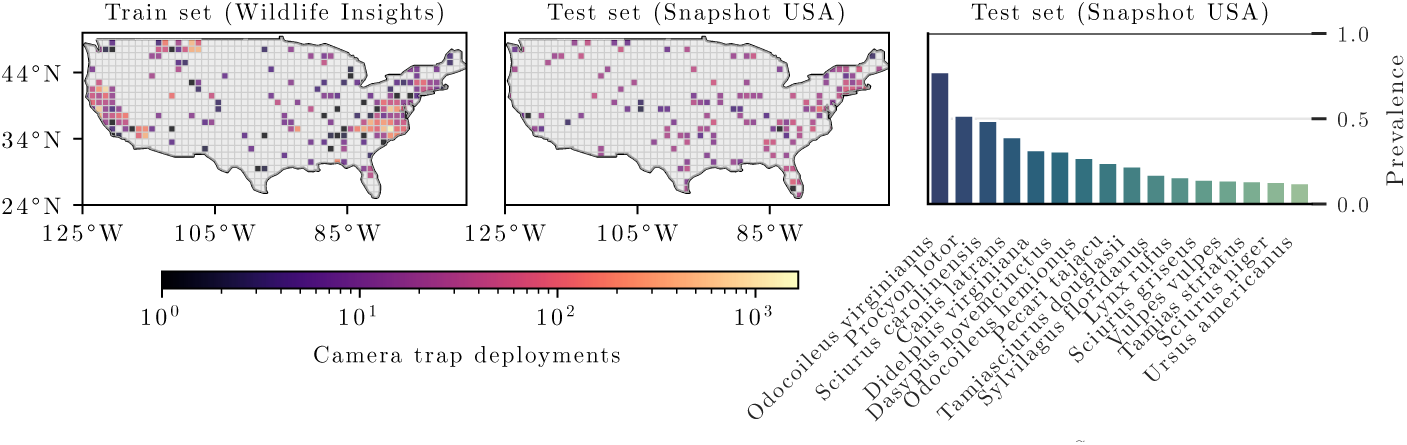
We train and evaluate occupancy models for 16 species of varying prevalence across the contiguous United States. The left and center panels show the density of train and test sites, respectively. The right panel shows prevalence across our focal species, here referring to the fraction of test-set sites within each species’ range (see Methods 4.1) where the species was detected at least once.

### 4.2 Occupancy model

To take advantage of the rich detection/non-detection data provided by camera trapping surveys, we leverage the occupancy model proposed by MacKenzie et al. [37]. Instead of assuming perfect observations of species presence or absence, the occupancy status of a species is modeled as a Bernoulli latent variable *z_i_*∈ {0, 1} with probability of occupancy *ψ_i_*, i.e.:

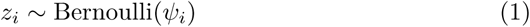

For each site *i* and discrete time steps *j* ∈ 1*, …, J_i_*, detection/non-detections *y_i,j_* ∈ {0, 1} are then modeled conditional on the occupancy status with probability of detection *p*. Specifically, we have:

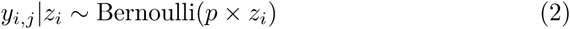

We model the probability of occupancy *ψ_i_* using a linear regression with the logit-link function (“logit-linear”), parameterized by coefficients *β* and predictors *X_i_* at site *i*:

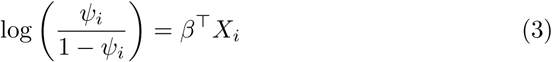

Although our model could also incorporate detection-level covariates, in this particular study, we forgo observation-level detection probability covariates and instead estimate a fixed detection probability *p* for each species across all sites to focus on how deep-learned features can improve occupancy-specific estimation. Incorporating detection covariates would add additional complexity that could make interpretation difficult in comparing candidate models.

As baseline predictors, we select a set of environmental variables commonly used for large-scale SDMs, consisting of bioclimatic variables [28] and soil properties [48]. We describe the full set of variables in Section C.1.

Additionally, we make use of satellite and ground-level imagery to capture environmental information about the location that is higher resolution and perhaps more nuanced than what is captured by the environmental variables described above. In order to do so, we make use of various machine learning models that were trained using representation learning approaches (see Section A.3).

For simplicity, we do not consider species interactions and fit separate models for each species. Since our occupancy model expects observations at discrete time-steps, but our data consists of detections / non-detections at continuous times, we discretize observations *y_i,j_* at site *i* over one-week-long time steps *j*. We discretize such that any week with at least one observation counts as observed detection (*y_i,j_* = 1) and any week without at least one observation counts as a non-detection (*y_i,j_*= 0). This type of discretization into week-long time-steps is in line with prior work [33, 44] and provides a sufficient number of repeated samples while keeping computational cost reasonable. We exclusively use human-labeled camera trap images to minimize the chance for AI-induced misclassifications.

To estimate the parameters of this hierarchical Bayesian model, we have to integrate or sample over the posterior distribution of the latent states. To achieve this, we introduce a new open-source implementation of occupancy models we call *Biolith*. Biolith is written in the Python programming language and uses the NumPyro probabilistic programming library [46] to perform Hamiltonian Monte Carlo inference using the No-U-Turn Sampler [29]. Being written in Python, our implementation enables machine learning to easily be incorporated into ecological modeling. More details on Biolith can be found in Section C.2.

To obtain representations of ground-level imagery, we sample a single blank (i.e. containing no animal) daylight image per location and extract a high-dimensional representation *X*_img_ using a machine learning model. Depending on the exact model used, these representations vary from hundreds to thousands of dimensions and thus lead to significant memory requirements. To alleviate this and keep all data in memory, we apply principal component analysis and keep only the 128 dimensions that explain the most variance. We follow the same process to extract representations of satellite imagery *X*_sat_. Finally, we simply concatenate all site-specific predictors into a single multimodal habitat representation *X_i_* for site *i*.

Reducing model complexity is crucial for reducing the risk of overfitting and for making models easier to interpret. Since our predictors *X* have on the order of 100 dimensions, fitting and performing model selection across all possible subsets of predictors is infeasible. Instead, we exclusively rely on regularization to reduce model complexity, a common approach, for example, in MaxEnt SDMs [64]. We impose L2 or L1 regularization by placing Gaussian or Laplace priors on the model coefficients, respectively [62]. To find the optimal configuration of prior type and scale for each species and combination of modalities, we perform 3-fold cross validation on the train set, stratified by naive occupancy (i.e., whether the site has at least one observation). This stratification ensures that there are sites with and without detections in the splits used to fit and validate the model. We then select the best configuration according to LPPD on the held-out portion, and re-fit on the entire train set.

### 4.3 Evaluation protocol

Since occupancy models assume there are no single ground truth species presence/absence labels (see Section A.2), only detections / non-detections, we evaluate using the likelihood of test-set detection / non-detection data under the fitted model through the posterior predictive distribution. To construct the likelihood measure for each candidate occupancy model, we draw *Q* posterior samples of parameters *θ*^(*q*)^ = {*β*^(*q*)^*, p*^(*q*)^} and summarize across posterior draws using the log point-wise predictive density (LPPD) [22]:

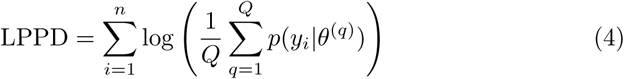

where *n* is the number of sites, *J_i_*is the number of visits at site *i*, *y_i_*is the observed detection history for site *i* across *J_i_* revisits, *y_ij_* is the observed detection at site *i* and visit *j*, and *p*^(*q*)^ is the detection probability in sample *q*.

Intuitively, the LPPD provides a measure of how divergent a set of observed data is from the posterior distribution of a fitted model. It is helpful because it allows us to evaluate model fit using only repeated observations and without having explicit ground truth labels for species presence/absence. However, similar to other likelihood-based measures, the magnitude of LPPD can be difficult to interpret and compare across species. To create a metric which provides more insight into the absolute performance of a given model and make it easier to compare across species, we compute a linear transformation of LPPD that places each model on a 0-1 scale defined by the LPPD of a “null” and an “oracle” model within a given species. The null model is a model that incorporates nothing but an intercept term for occupancy and detection, and is therefore limited to estimating a single mean occupancy probability over all sites. In contrast, the oracle model is fit to the otherwise held-out *test* data using site-level random effects. The oracle model therefore acts as an upper bound and intuitively represents a model whose predictors and parameters perfectly mirror the underlying ecological occurrence process on the test data. Specifically, given the LPPD of null and oracle models LPPD_null_, LPPD_oracle_, we normalize a given LPPD as follows:

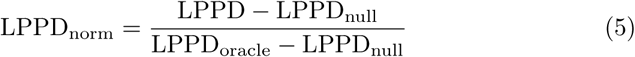

We note that this normalization can result in negative values if the model of interest fails to outperform the null model.

#### 4.3.1 Statistical Evaluation

We evaluate individual combinations of predictors using paired Wilcoxon signed-rank tests between the *X*_env_ + *X*_sat_ + *X*_img_ model and all other predictor combinations (one-tailed *H*_1_: 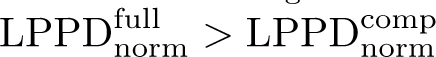; Holm correction across these pairwise comparisons). Additionally, we perform one-sample Wilcoxon signed-rank tests between all combinations of predictors and the null model (one-tailed *H*_1_: LPPD_norm_ *>* 0), and a three-way analysis of variance (ANOVA) using all predictor combinations and considering each of *X*_env_, *X*_sat_, and *X*_img_ as fixed factors, with Tukey HSD post-hoc corrections where applicable.

### 4.4 Describing habitat elements in natural language

To enable interpreting and simplifying complex occupancy models, we propose the technique illustrated in Figure 3. We first use the coefficients corresponding to ground-level image features *β*_img_ and compute 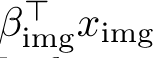 to rank images from most to least likely occupied by a given species. We then apply an adaptation of VisDiff [16] to describe habitat elements in natural language that are present in likely occupied images but absent from likely unoccupied images.

VisDiff uses three different types of models: contrastively trained vision-language models, image-conditional text generation models, and general large language models (LLMs).

**Contrastively trained vision-language models** such as CLIP [50] are trained to map images and corresponding natural language captions to similar embeddings in a so-called shared embedding space, and non-corresponding image-caption pairs to dissimilar embeddings. These models can be used for measuring the alignment between an image *I* and caption *C*, by computing the embeddings for image *z_I_*and caption *z_C_*, and then computing the cosine similarity between both embeddings: 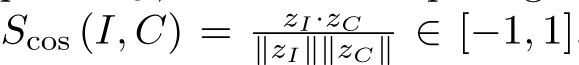. For example, an image *I* depicting a dog and a caption *C* saying “a dog sitting down” would have higher similarity *S*_cos_ (*I, C*) than an image of a cat with the same caption *C*.

**Image-conditional text generation models** are models that *generate* captions based on an input image. They are particularly useful for our use-case since they allow generating open-ended captions, as opposed to contrastively trained vision-language models, which require upfront hypotheses. Here we use the BLIP-2 [36].

**Large language models or LLMs** are models that take variable-length text (or sometimes other modalities) as input and produce variable-length output, excelling at general problem solving tasks.

#### We now leverage these different types of models using an adaptation of VisDiff [16]

We start by captioning 20 of the most and least likely occupied images, respectively, using the BLIP-2 model [36]. This results in captions such as “An image of a calm deciduous forest scene with rays of light shining through the forest canopy”. We then query GPT-5.2 to describe the differences between the captions of likely occupied vs. likely unoccupied images, resulting in a set of hypotheses. Finally, we use CLIP [50] to rank hypotheses according to how well they separate likely occupied from likely unoccupied images, keeping only the top five hypotheses. We note that, while the initial captions generated by BLIP-2 contain a high amount of irrelevant information, the two subsequent steps tend to focus the hypotheses onto the relevant ecological information.

## Code availability

Custom code used to generate the results reported in this study, together with the trained model weights and scripts needed to reproduce the experiments, is publicly available on GitHub at https://github.com/timmh/mmocc. The ecological modeling framework Biolith used in this work is publicly available on GitHub at https://github.com/timmh/biolith.

## Acknowledgments

This work was supported in part by the MIT-IBM Watson AI Lab, the MIT METEOR Program, the AI and Biodiversity Change (ABC) Global Center, which is funded by the US National Science Foundation under Award No. 2330423 and Natural Sciences and Engineering Research Council of Canada under Award No. 585136, NSF CAREER Award No. 2441060, the MIT-Google program for Computing Innovation and Planetary Health, the Abdul Latif Jameel Water and Food Systems Lab at MIT, and Goldman Sachs. This work draws on research supported in part by the Social Sciences and Humanities Research Council. We thank Bill McShea, Roland Kays, and all other collaborators of Snapshot USA.

## Wildlife Insights data references

A One Tam project managed collaboratively by the National Park Service, California State Parks, Marin Water, Marin County Parks and Golden Gate National Parks Conservancy. Last updated august 2025. http://n2t.net/ark:/63614/w12001631, 2017. Accessed via wildlifeinsights.org on September 1, 2025.

A One Tam project managed collaboratively by the National Park Service, California State Parks, Marin Water, Marin County Parks and Golden Gate National Parks Conservancy. Last updated july 2025. http://n2t.net/ark:/63614/w12001634, 2017. Accessed via wildlifeinsights.org on September 1, 2025.

A One Tam project managed collaboratively by the National Park Service, California State Parks, Marin Water, Marin County Parks and Golden Gate National Parks Conservancy. Last updated august 2025. http://n2t.net/ark:/63614/w12001639, 2017. Accessed via wildlifeinsights.org on September 1, 2025.

A One Tam project managed collaboratively by the National Park Service, California State Parks, Marin Water, Marin County Parks and Golden Gate National Parks Conservancy. Last updated august 2025. http://n2t.net/ark:/63614/w12001648, 2017. Accessed via wildlifeinsights.org on September 1, 2025.

Aiello, C. M., Galloway, N. L., Fratella, K., Prentice, P. R., Darby, N. W., Hughson D., Epps, C. W. Last updated november 2024. http://n2t.net/ark:/63614/w12003007, 2018. Accessed via wildlifeinsights.org on September 1, 2025.

Bresnan, C. Last updated november 2024. http://n2t.net/ark:/63614/w12006261, 2023. Accessed via wildlifeinsights.org on September 1, 2025.

Bresnan, C., Shamon, H. Last updated september 2024. http://n2t.net/ark:/63614/w12004234, 2022. Accessed via wildlifeinsights.org on September 1, 2025.

Casady, D. Last updated march 2025. http://n2t.net/ark:/63614/w12002889, 2016. Accessed via wildlifeinsights.org on September 1, 2025.

Casady, D. Last updated may 2025. http://n2t.net/ark:/63614/w12002877, 2017. Accessed via wildlifeinsights.org on September 1, 2025.

Casady, D. Last updated march 2025. http://n2t.net/ark:/63614/w12002923, 2017. Accessed via wildlifeinsights.org on September 1, 2025.

Casady, D. Last updated january 2025. http://n2t.net/ark:/63614/w12002901, 2019. Accessed via wildlifeinsights.org on September 1, 2025.

Critter Camera Data. http://n2t.net/ark:/63614/w12001313. Accessed via wildlifeinsights.org on September 1, 2025.

Davis, J. Last updated august 2025. http://n2t.net/ark:/63614/w12004572, 2016. Accessed via wildlifeinsights.org on September 1, 2025.

Dorcy, J., Rich, L., Furnas, B., Dorcy-Ponce, J. Last updated march 2025. http://n2t.net/ark:/63614/w12002828, 2017. Accessed via wildlifeinsights.org on September 1, 2025.

Flores, N., singh, a., Rich, L., Cook, V., Vollmer, J., Whetzel, M. Last updated july 2025. http://n2t.net/ark:/63614/w12004221, 2019. Accessed via wildlifeinsights.org on September 1, 2025.

Flores, N., Smithsonian, e., Kays, R. Last updated march 2024. http://n2t.net/ark:/63614/w12003860, 2008. Accessed via wildlifeinsights.org on September 1, 2025.

Flores, N., Smithsonian, e., Kays, R., Chrysafis, P. Last updated april 2023. http://n2t.net/ark:/63614/w12003854, 2011. Accessed via wildlifeinsights.org on September 1, 2025.

Flores, N., Smithsonian, e., McShea, W., Shamon, H. Last updated august 2025. http://n2t.net/ark:/63614/w12006132, 2013. Accessed via wildlifeinsights.org on September 1, 2025.

Forrester, T. . Last updated january 2023. http://n2t.net/ark:/63614/w12004287, 2011. Accessed via wildlifeinsights.org on September 1, 2025.

Forrester, T. . Last updated april 2023. http://n2t.net/ark:/63614/w12004316, 2011. Accessed via wildlifeinsights.org on September 1, 2025.

Forrester, T. Last updated august 2025. http://n2t.net/ark:/63614/w12004302, 2000. Accessed via wildlifeinsights.org on September 1, 2025.

Green, A. Last updated june 2025. http://n2t.net/ark:/63614/w12003225, 2020. Accessed via wildlifeinsights.org on September 1, 2025.

C. P. Hansen, A. W. Parsons, R. Kays, and J. J. Millspaugh. Does use of backyard resources explain the abundance of urban wildlife? *Frontiers in Ecology and Evolution*, 8:570771, 2020. URL http://n2t.net/ark:/63614/w12004526. Accessed via wildlifeinsights.org on September 1, 2025.

Hatfield, B., Rich, L., Malleshappa, V. Last updated august 2025. http://n2t.net/ark:/63614/w12002818, 2020. Accessed via wildlifeinsights.org on September 1, 2025.

Higdon, S. Last updated april 2023. http://n2t.net/ark:/63614/w12004592, 2000. Accessed via wildlifeinsights.org on September 1, 2025.

Jachowski, D.S.; Katzner, T.E.; Harris, S.N.; Marneweck, C.J. Last updated august 2023. http://n2t.net/ark:/63614/w12001570, 2007. Accessed via wildlifeinsights.org on September 1, 2025.

Jachowski, D.S.; Katzner, T.E.; Harris, S.N.; Marneweck, C.J. Last updated july 2024. http://n2t.net/ark:/63614/w12003589, 2010. Accessed via wildlifeinsights.org on September 1, 2025.

Jansen, P. Last updated february 2023. http://n2t.net/ark:/63614/w12004587, 2015. Accessed via wildlifeinsights.org on September 1, 2025.

Johnson, Aaron P., Davis, Allegra G., Garcia, Diego G., Tobio, Ana, Wesling, Lauren A., Dibble, Josh. . Last updated july 2025. http://n2t.net/ark:/63614/w12004413, 2015. Accessed via wildlifeinsights.org on September 1, 2025.

Jones, A. Last updated september 2024. http://n2t.net/ark:/63614/w12003849, 2016. Accessed via wildlifeinsights.org on September 1, 2025.

Kalies, E.L. Last updated november 2022. http://n2t.net/ark:/63614/w12004569, 2016. Accessed via wildlifeinsights.org on September 1, 2025.

Kays, R. . Last updated june 2025. http://n2t.net/ark:/63614/w12004713, 2000. Accessed via wildlifeinsights.org on September 1, 2025.

Kays, R. . Last updated january 2023. http://n2t.net/ark:/63614/w12004780, 2011. Accessed via wildlifeinsights.org on September 1, 2025.

Kays, R. . Last updated september 2024. http://n2t.net/ark:/63614/w12004748, 2012. Accessed via wildlifeinsights.org on September 1, 2025.

Kays, R. . Last updated june 2025. http://n2t.net/ark:/63614/w12004318, 2015. Accessed via wildlifeinsights.org on September 1, 2025.

Kays, R. . Last updated march 2025. http://n2t.net/ark:/63614/w12004782, 2017. Accessed via wildlifeinsights.org on September 1, 2025.

Kays, R., Levin, M., Kalies, L. Last updated february 2023. http://n2t.net/ark:/63614/w12004578, 2022. Accessed via wildlifeinsights.org on September 1, 2025.

Kays, R., Support, W., Ahumada, J., Alyetama, M. Last updated august 2025. http://n2t.net/ark:/63614/w12003602, 2021. Accessed via wildlifeinsights.org on September 1, 2025.

Kays, Roland; Hess, George . Last updated august 2025. http://n2t.net/ark:/63614/w12004290, 2000. Accessed via wildlifeinsights.org on September 1, 2025.

kumar, s., Stutzman, J. Last updated may 2024. http://n2t.net/ark:/63614/w12003608, 2021. Accessed via wildlifeinsights.org on September 1, 2025.

LaPoint, S. . Last updated november 2022. http://n2t.net/ark:/63614/w12004076, 2019. Accessed via wildlifeinsights.org on September 1, 2025.

Long, R. . Last updated january 2024. http://n2t.net/ark:/63614/w12004625, 2000. Accessed via wildlifeinsights.org on September 1, 2025.

M. Lasky, et al. CAROLINA CRITTERS: a collection of camera trap data from wildlife surveys across North Carolina. Ecology. e03372 (2021). Last updated december 2023. http://n2t.net/ark:/63614/w12004307, 2000. Accessed via wildlifeinsights.org on September 1, 2025.

Malleshappa, V., Flores, N., Rich, L., Found-Jackson, C., Carrothers, D., Anderson, S. Last updated march 2025. http://n2t.net/ark:/63614/w12003821, 2019. Accessed via wildlifeinsights.org on September 1, 2025.

Malleshappa, V., Smithsonian, e., Kays, R. Last updated july 2023. http://n2t.net/ark:/63614/w12004600, 2000. Accessed via wildlifeinsights.org on September 1, 2025.

Malleshappa, V., Smithsonian, e., McShea, W., Kays, R. Last updated june 2023. http://n2t.net/ark:/63614/w12004286, 2013. Accessed via wildlifeinsights.org on September 1, 2025.

Malleshappa, V., Smithsonian, e., McShea, W., Schuttler, S., Kays, R. Last updated april 2023. http://n2t.net/ark:/63614/w12004293, 2014. Accessed via wildlifeinsights.org on September 1, 2025.

Malleshappa, V., Smithsonian, e., Rooney, B., Kays, R. Last updated july 2025. http://n2t.net/ark:/63614/w12004090, 2012. Accessed via wildlifeinsights.org on September 1, 2025.

Malleshappa, V., Smithsonian, e., Schuttler, S., Kays, R. Last updated december 2023. http://n2t.net/ark:/63614/w12004311, 2014. Accessed via wildlifeinsights.org on September 1, 2025.

Martin, Marie E., Green, David S., and Matthews, Sean M. http://n2t.net/ark:/63614/w12001294. Accessed via wildlifeinsights.org on September 1, 2025.

S. McMurry, A. W. Parsons, M. Lasky, H. Boone, R. Kays, M. Snider, E. Kays, J. Luongo, J. Clark, C. L. Scher, and W. J. McShea. Standardized camera trapping data from National Ecological Observatory Network (NEON) sites and other forest plots. Ecology, in review. Accessed via wildlifeinsights.org on September 1, 2025. URL http://n2t.net/ark:/63614/w12004578.

McMurry, S., Kays, R., Alyetama, M., Patel, L. Last updated june 2025. http://n2t.net/ark:/63614/w12006449, 2023. Accessed via wildlifeinsights.org on September 1, 2025.

McMurry, S., Kays, R., Spurlin, J., Martin, G., Frech, G., Barajas-Salazar, K., Snider, M. Last updated october 2023. http://n2t.net/ark:/63614/w12004251, 2022. Accessed via wildlifeinsights.org on September 1, 2025.

McShea, W. . Last updated june 2025. http://n2t.net/ark:/63614/w12004765, 1999. Accessed via wildlifeinsights.org on September 1, 2025.

McShea, W. . Last updated august 2025. http://n2t.net/ark:/63614/w12004781, 2000. Accessed via wildlifeinsights.org on September 1, 2025.

Mueldener, M. Last updated april 2023. http://n2t.net/ark:/63614/w12004588, 2017. Accessed via wildlifeinsights.org on September 1, 2025.

Myers, J. . Last updated april 2023. http://n2t.net/ark:/63614/w12004295, 2014. Accessed via wildlifeinsights.org on September 1, 2025.

Nelson, D. L. Shamon, H. Jachowski, D. S. Last updated april 2025. http://n2t.net/ark:/63614/w12003597, 2021. Accessed via wildlifeinsights.org on September 1, 2025.

Nelson, D. L. Shamon, H. Jachowski, D. S. Last updated november 2024. http://n2t.net/ark:/63614/w12004526, 2022. Accessed via wildlifeinsights.org on September 1, 2025.

Nelson, Dana L. . Last updated july 2025. http://n2t.net/ark:/63614/w12002979, 2021. Accessed via wildlifeinsights.org on September 1, 2025.

Rich, L. Last updated april 2025. http://n2t.net/ark:/63614/w12003401, 2012. Accessed via wildlifeinsights.org on September 1, 2025.

Rich, L. N., Toenies, M., Cornelius, N., Boynton, M., Chappell, E. Last updated august 2025. http://n2t.net/ark:/63614/w12001650, 2022. Accessed via wildlifeinsights.org on September 1, 2025.

Rooney, Brigit; McShea, William. Last updated august 2025. http://n2t.net/ark:/63614/w12006220, 2023. Accessed via wildlifeinsights.org on September 1, 2025.

Rota, C. . Last updated november 2023. http://n2t.net/ark:/63614/w12003869, 2016. Accessed via wildlifeinsights.org on September 1, 2025.

Rota, C. Last updated december 2023. http://n2t.net/ark:/63614/w12003912, 2016. Accessed via wildlifeinsights.org on September 1, 2025.

Samsoe, E. Last updated october 2023. http://n2t.net/ark:/63614/w12003109, 2007. Accessed via wildlifeinsights.org on September 1, 2025.

Sargis, E. , Widness, J. Last updated september 2024. http://n2t.net/ark:/63614/w12003858, 2017. Accessed via wildlifeinsights.org on September 1, 2025.

Sevin, J. Last updated august 2025. http://n2t.net/ark:/63614/w12004615, 2020. Accessed via wildlifeinsights.org on September 1, 2025.

Shamon, H. Last updated may 2023. http://n2t.net/ark:/63614/w12003074, 2021. Accessed via wildlifeinsights.org on September 1, 2025.

Shamon, H. Last updated july 2023. http://n2t.net/ark:/63614/w12006229, 2023. Accessed via wildlifeinsights.org on September 1, 2025.

singh, a., Malleshappa, V., Rich, L., Cook, V., Vollmer, J., Whetzel, M. Last updated july 2025. http://n2t.net/ark:/63614/w12003570, 2020. Accessed via wildlifeinsights.org on September 1, 2025.

Sink, C. Last updated june 2023. http://n2t.net/ark:/63614/w12002779, 2020. Accessed via wildlifeinsights.org on September 1, 2025.

Smithsonian, e. Last updated april 2023. http://n2t.net/ark:/63614/w12002684, 2020. Accessed via wildlifeinsights.org on September 1, 2025.

Smithsonian, e. Last updated april 2023. http://n2t.net/ark:/63614/w12002685, 2020. Accessed via wildlifeinsights.org on September 1, 2025.

Smithsonian, e., Farkas, D. Last updated december 2023. http://n2t.net/ark:/63614/w12004582, 2011. Accessed via wildlifeinsights.org on September 1, 2025.

Smithsonian, e., McShea, W. Last updated march 2025. http://n2t.net/ark:/63614/w12003833, 2007. Accessed via wildlifeinsights.org on September 1, 2025.

Smithsonian, e., McShea, W. Last updated april 2023. http://n2t.net/ark:/63614/w12004583, 2017. Accessed via wildlifeinsights.org on September 1, 2025.

Smithsonian, e., McShea, W., Shamon, H. Last updated june 2025. http://n2t.net/ark:/63614/w12006140, 2012. Accessed via wildlifeinsights.org on September 1, 2025.

Tucker, J., Support, B., Heiman, J., Heiman, J., Eyes, S. Last updated july 2025. http://n2t.net/ark:/63614/w12004223, 2021. Accessed via wildlifeinsights.org on September 1, 2025.

USDA Forest Service, Pacific Southwest Region, Carnivore Monitoring Program. http://n2t.net/ark:/63614/w12000517. Accessed via wildlifeinsights.org on September 1, 2025.

VanNiel, J. Last updated april 2023. http://n2t.net/ark:/63614/w12003830, 2017. Accessed via wildlifeinsights.org on September 1, 2025.

Wainstein, M. Last updated april 2023. http://n2t.net/ark:/63614/w12003862, 2014. Accessed via wildlifeinsights.org on September 1, 2025.

Woodrow, A. Last updated october 2022. http://n2t.net/ark:/63614/w12003864, 2011. Accessed via wildlifeinsights.org on September 1, 2025.

## A Related work

### A.1 Species distribution modeling

In 1904, Grinnell first established qualitatively that patterns of species occurrence can be inferred from environmental variables such as humidity [85]. In the century since, a rich body of quantitative research has emerged around associating environmental variables with the occurrence of specific species. The most widespread approaches are called species distribution models (SDMs), which generally aim to model the relationship between species and habitats through environmental variables such as climate, elevation, or human presence. These models are used by land managers and policy makers to, e.g., understand risks to species and prioritize conservation action, critical for both ecological research and practical on-the-ground species and habitat conservation. The model methodologies used for SDMs and the interpretation of the model results are primarily driven by what kind of species occurrence records are used as inputs. There are two primary types of data commonly used to fit SDMs: (1) presence-only data and (2) detection/non-detection data.

We note that detection/non-detection data and presence/absence are often used interchangeably in the literature to express observations where there can be either positives or negatives for a species occurrence at a location. We use the language of detection/non-detection to describe the data because the treatment of non-detections as true absences can bias or confound SDM estimation [86, 99, 100]. Using the terms presence-absence implies that the absence data is reflective of true “non-occupancy” and not a lack of detection.

The majority of SDM methods are designed to address the specific nuances of either presence-only or detection/non-detection data, however there are some models that are designed to integrate both types [79, 78] or other types of occurrence records such as abundance data [95].

*Presence-only data*, in which only records that contain a positive detection or incidence of a species occurrence, is the most abundant data available for modeling. Due to a lack of “negative” detections in the data, methods developed for presence-only data typically derive either “pseudo-absence” or “background” locations [67]. For single-species distribution models, common model methodologies include generalized linear models, generalized additive models, MaxEnt [107], and random forest/tree-based models [123] and leverage environmental features to characterize the species distribution. The typical interpretation of the output of these models is not occupancy but rather “habitat suitability” – where locations more similar to sites where species have been detected in the environmental feature space are predicted to have higher values.

Over the last decade, neural-network-based SDMs have emerged that can scalably model presence-only species detection data through species co-occurrence [72], where many recent innovations focus on deep-learning primarily through either novel loss functions that have different mechanisms for defining and weighting pseudo-absences and/or the inclusion of remote sensing features [124, 84, 75]. For example, Cole et al. have shown that spatial implicit neural representations (SINR) are effective at learning from large-scale citizen science data across hundreds of thousands of species and perform well for tasks like range estimation from jointly modeling presence-only species occurrence records with few or no environmental characteristics [75]. Heterogeneous graph neural networks have also emerged as deep-learning approaches that can add additional structure in learning species distributions and interactions from presence-only data [88].

*Detection/non-detection data* contains records of both positive species detections, where target species were found at surveyed locations, as well as non-detections across different places or areas for a given survey – where non-detections indicate a species that was included in the target species set of a survey, but not found at the location during the survey time. Detection/non-detection data can be used as positives/negatives for training binary classification models, but these results can be highly misleading, similar to results from presence-only data [86]. Because detection is frequently imperfect, a species can still be present or occupy a given habitat without being detected, which means that *observed* non-detection (or absence) does not equal *true* absence [99].

The probability of detecting a species, if the species is truly present, may be highly sensitive to the observation method (e.g., human experts or automated sensors such as camera traps) and survey effort (e.g., survey duration, time of day, area surveyed, and number of repeated visits, if any). Accounting for these effects is difficult, especially if observer effects and effort are unknown or not characterized in a standardized way across surveys, as is the case with large-scale citizen science data repositories. Even if there is a combination of presence-only data and detection/non-detection data, it is common for many SDMs to reduce their training data to presence-only records because dealing with the nuances of survey details across many study design types increases the complexity. The focus of this paper is leveraging large-scale detection/non-detection records from repeated surveys using wildlife camera trap image data, but could be extended for integrated models that include both presence-only and detection/non-detection surveys.

### A.2 Occupancy modeling

In contrast to presence-only SDMs, occupancy models disentangle the probability of species presence (or *occupancy*) from the probability of detecting said species [101]. As opposed to treating a species’ use of a habitat as a binary classification model on detection/non-detection status, occupancy models treat the habitat usage or site occupancy as a latent, or unobserved variable, where there is no observable “ground truth” absence (or presence if false-positives are assumed). Occupancy models are able to estimate separate processes for the probability of detection and probability of occupancy by incorporating repeated survey data of species detection and non-detection - where the rate of positive detections versus non-detections allows for estimation of the probability of detecting a species given a site is occupied. Site occupancy is then estimated through a posterior distribution given the observed data, which could also include covariates for both the occupancy and the detection processes.

The ability to account for separate detection versus occupancy processes is particularly crucial for animals that are difficult to detect, and for which naively assuming absence based on few negative observations would significantly bias the occupancy estimate to a lower than actual level. For example, consider a nocturnal and elusive species like the American marten (Martes americana). An expert surveying this species during the day could be standing right on top of a marten den while at the same time being unaware of their presence, even over the course of multiple visits. In practice, not accounting for imperfect detection of species like the American marten can lead to an underestimation in presence at 15–52% of surveyed sites [115].

Occupancy models have been widely applied to infer species distributions, population trends, and responses to environmental variables. Early applications often relied on manual surveys (e.g., visual observations by researchers in the field) and were thus limited to relatively small areas. In recent years, improved autonomous sensor technology such as camera traps and autonomous acoustic recording units has greatly expanded the scale and efficiency of occupancy studies.

Just like other SDMs, occupancy models typically use a small set of environmental variables as predictors of occupancy. Common environmental variables range from habitat structure (e.g., land-cover heterogeneity [103] and vegetation structure [98]), topography (e.g., elevation [66]), climate (e.g., mean annual temperature [66]), and human influence (e.g., presence of non-native fish species [87]). Other works are simply concerned with estimating average occupancy and thus forgo environmental variables as occupancy predictors and estimate a constant occupancy probability across sites [96].

### A.3 Representation learning

Deep learning models are known to learn powerful, high-level representations of their input data. Here we describe various ways in which these representations are learned.

#### Supervised learning approaches can learn useful representations from manually labeled data

For example, an image classification model such as SpeciesNet [83], which is trained to recognize various animal species, will likely learn representations associated with the species’ specific shapes and textures. For intuition about what type of information these types of representations capture, we refer the reader to the work of [125].

#### Alternatively, representations can be learned without manually labeled data using a process called self-supervised learning (SSL)

Here, the model uses structure within the training data to obtain useful learning signals. For example, approaches such as MoCo [89] and SimCLR [73] process two slightly altered versions of a single image (e.g., different cropped regions) and try to make their representations as similar as possible, thereby capturing the high-level information common across both altered versions while minimizing the impact of “nuisance” information that is unique to each individual version and therefore the result of the alterations (contrastive learning). These types of approaches have since been improved upon primarily by scaling the volume of training data, such as by DINOv2, which is trained on 1.2B images [104]. Although popularized by machine learning models trained on regular imagery, SSL methods have furthermore successfully been applied to learn generally useful representations of single-time-step satellite imagery [94, 102, 65]. More recent methods incorporate temporal information by focusing on series of satellite images over time, thereby capturing effects such as seasonality [121, 122, 93, 118, 70].

### A.4 Deep features in ecological models

There has been growing interest in the research community in combining raw satellite imagery with features extracted from deep learning models as predictors for SDMs [74, 76, 119]. [120] finds that leveraging satellite imagery in the RGB and NIR bands alongside environmental variables improves prediction of bird species encounter rates. Similarly, Sat-SINR [77] extends SINR [75] (see section A.1) by adding satellite imagery as additional predictor, alongside environmental variables and geographic location, to map the distribution of plant species at high resolution.

CRISP [92], BirdSAT [112] learn useful representations of satellite imagery for species classification and distribution modeling by contrastively training the model to match satellite and ground-level images taken from the same location (see section A.3). This objective encourages the model to emphasize whatever information is consistent across both views. Because they use ground-level imagery only during training, these methods can fundamentally make predictions anywhere on Earth using satellite imagery alone. However, much of the information visible in ground-level photos is not observable from space, which inherently limits how informative the learned satellite representations can be. In contrast to these approaches, our method can exploit the full richness of ground-level imagery at inference time.

TaxaBind [113] likewise employs a contrastive training objective across ground-level and satellite imagery, audio, geographic location, environmental variables and taxonomic hierarchy, and employs an optional multimodal patching technique that retains some unique information from different modalities. Among other downstream tasks, TaxaBind explores species distribution modeling qualitatively by comparing generated maps with citizen science observations. In contrast, here we are focusing on Bayesian hierarchical occupancy models that explicitly account for imperfect detection, and we evaluate those models quantitatively.

## B Additional results

In this section we describe additional experiments and corresponding results. Supplemental Table B.1 shows the full predictive performance across all species and combinations of modalities that underlies the results in fig. 2. Section B.1 evaluates non-hierarchical models that do not explicitly model detection probability while section B.2 evaluates alternative deep feature extractors. Section B.3 contains the full list of natural-language habitat elements.

**Table B.1:**
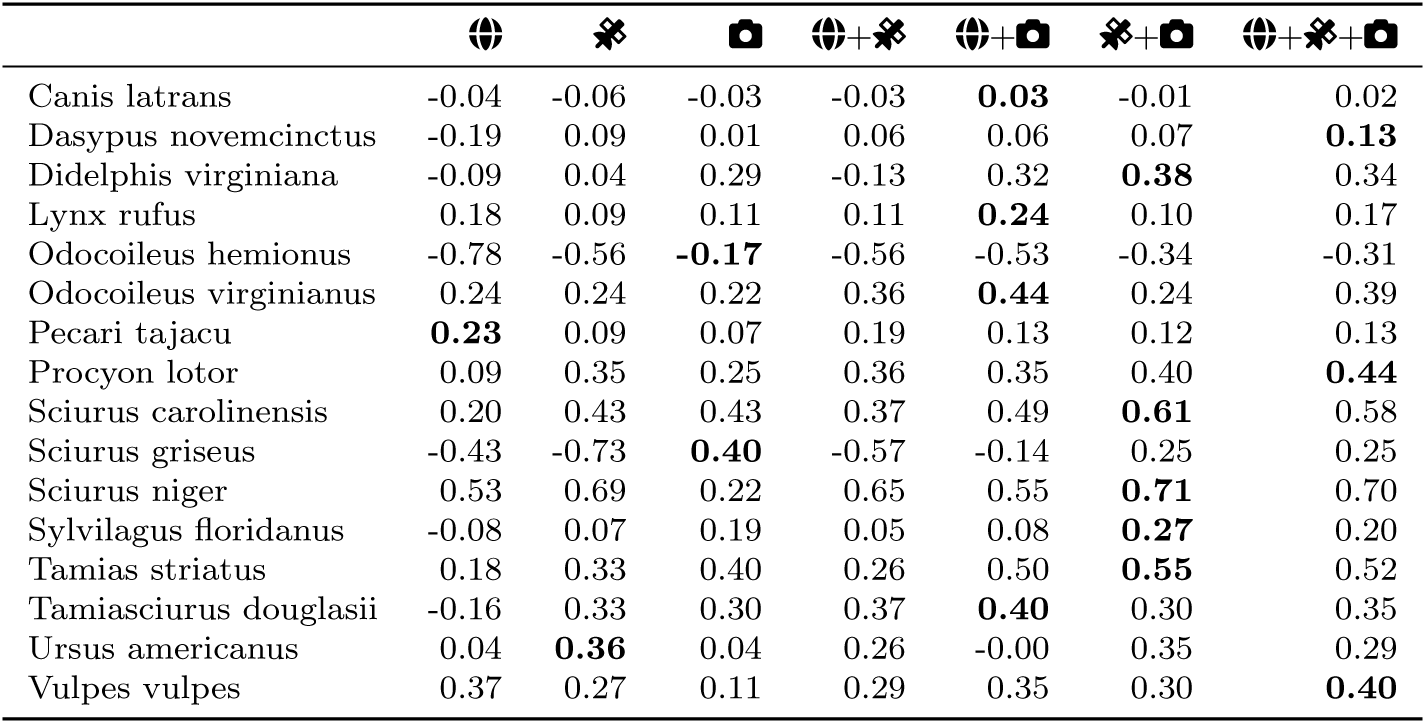
Test-set predictive performance as measured by normalized LPPD for different combinations of environmental variables 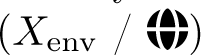, satellite imagery embeddings 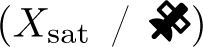, and ground-level imagery embeddings 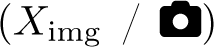. The best-performing model per species is bolded. **Overall, incorporating additional modalities improves predictive performance, with most species being modeled best when including ground-level imagery.** Entries outperforming the null model, which estimates a constant occupancy probability across all sites, are positive, others negative.

### B.1 Naive occupancy experiments

In addition to fitting full hierarchical occupancy models, we can fit a simpler class of models we call *naive* occupancy models. Instead of detections and non-detections, these models predict a single binary *occupied* or *unoccupied* class for each site. We say a site *i* is naively occupied 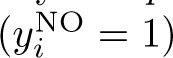 if there is at least one observation of the focal species at the site, otherwise we say the site is naively unoccupied 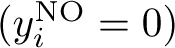.

While this simplified formulation does not properly account for imperfect detection, this might be of less concern for highly detectable species. Supplemental Figure B.1 shows that detection probability estimates for some of our focal species are relatively high. This simplified formulation then allows us to use any binary classification model such as logistic regression that can be optimized without having to rely on computationally expensive Markov-chain Monte Carlo. Additionally, naive occupancy can be evaluated using intuitive binary classification metrics such as average precision, also called PR-AUC. PR-AUC is the area under the precision-recall curve plotted over different confidence thresholds. It is popular for evaluating species distribution models, as it is robust to class imbalance, as, for example, induced by rare species [116]. Given presence-absence confidence scores *s_i_*produced by the naive occupancy classifier for site *i* and a set of confidence thresholds *t_n_*, recall and precision are computed for each of those thresholds as:

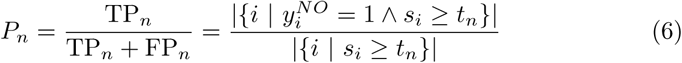

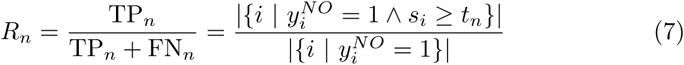

We then compute the approximate area under the precision-recall curve as:

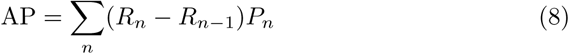

Supplemental Figure B.2 shows the improvement in PR-AUC of the best-performing ground-level image model over the best-performing non-ground-level image model and Supplemental Figure B.3 shows the same for satellite image features. Overall, we observe the same trends between naive and full occupancy models. Species benefiting more strongly from ground-level and satellite imagery in full occupancy models benefit similarly in naive occupancy models, and vice-versa.

**Figure B.1:**
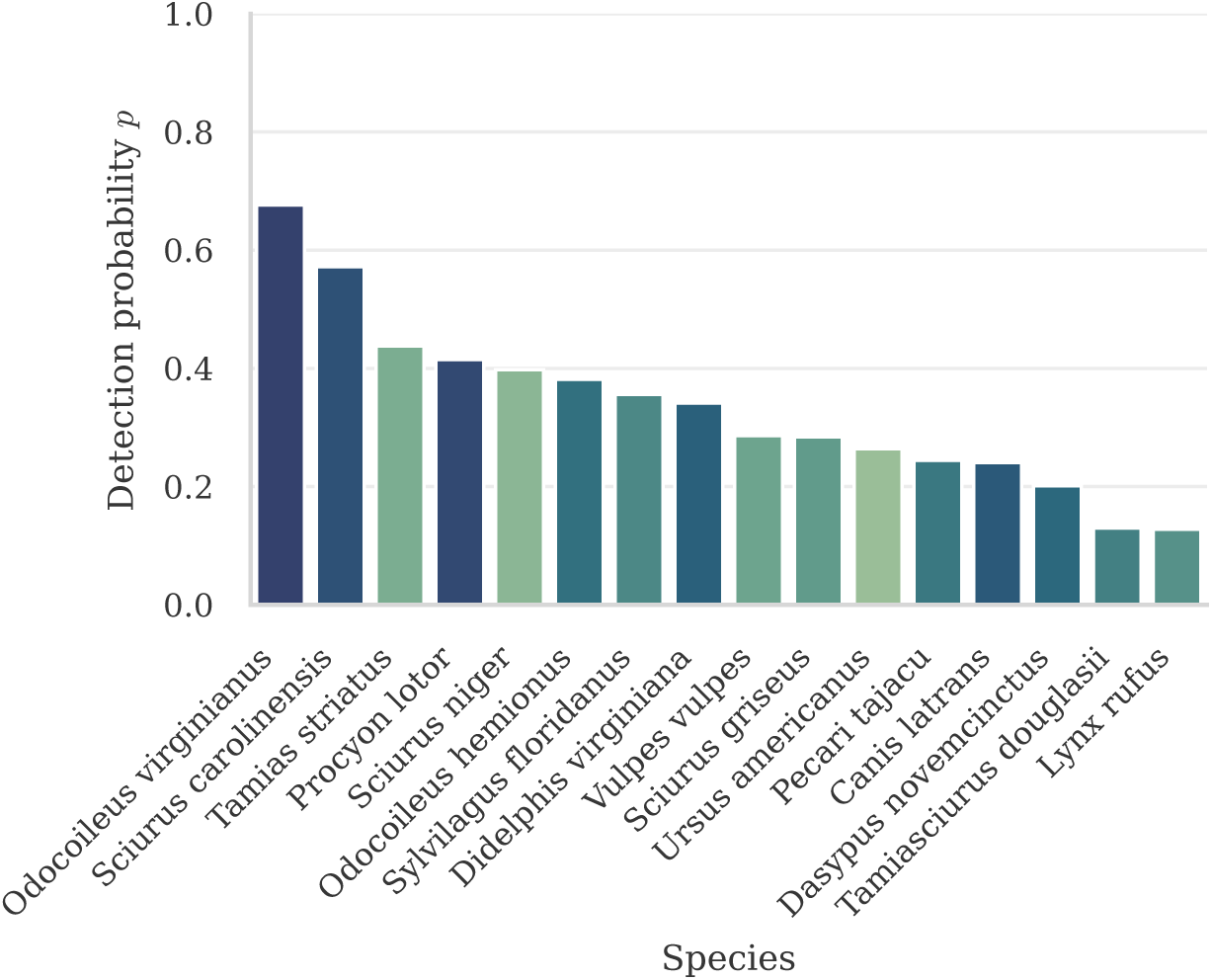
Posterior mean detection probability across species. **Detection probabilities are relatively high for many species, indicating that non-hierarchical logistic regression might be sufficient in some cases.** Blue colors represent more prevalent, green colors less prevalent species (see Methods 4.1).

**Figure B.2:**
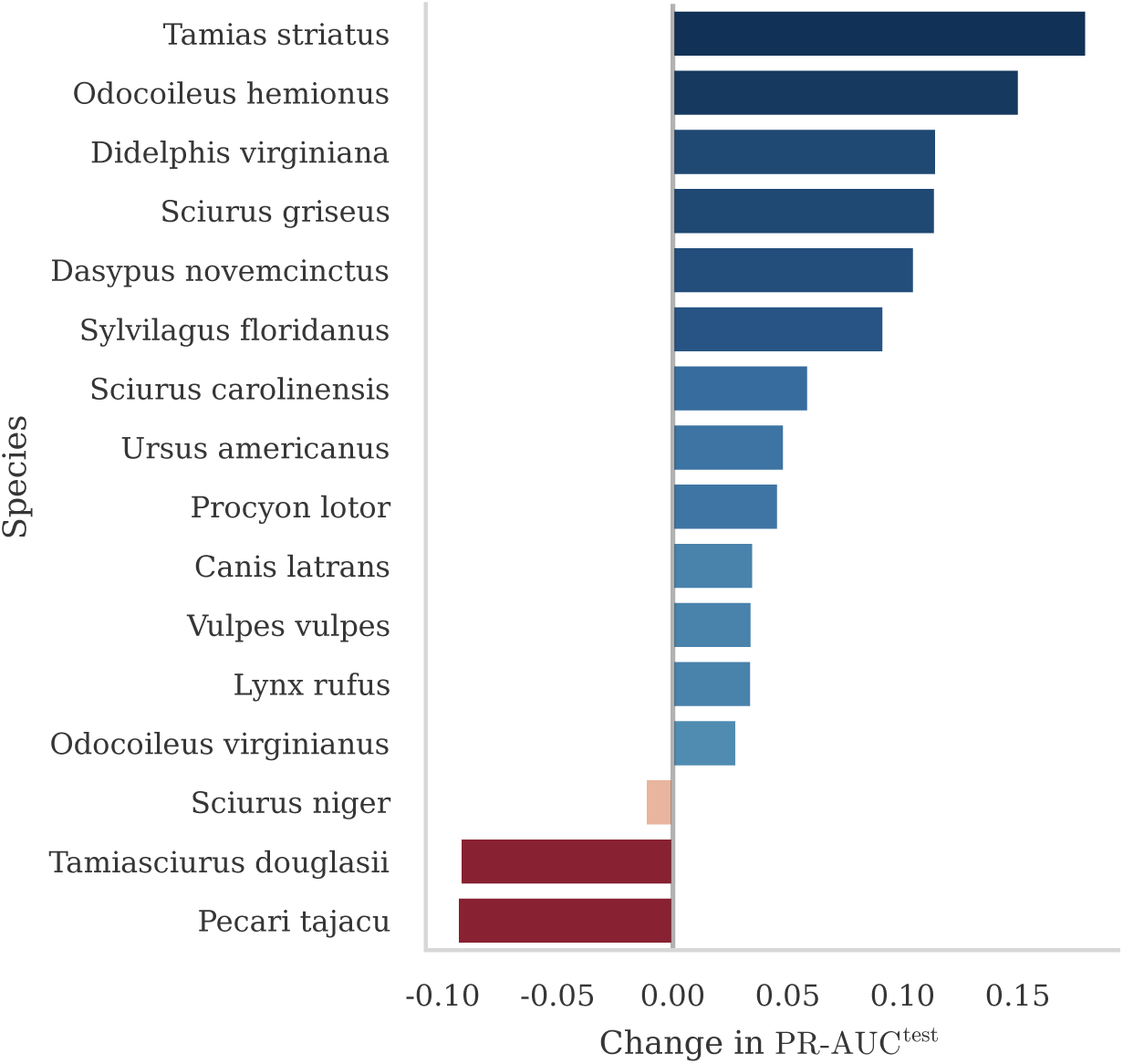
Test-set PR-AUC improvement of the best-performing model including ground-level image features over the best-performing model not including ground-level image features.

### B.2 Alternative deep feature extractors

In addition to DINOv2 [104] for ground-level images and the AlphaEarth Foundations model [70], we evaluate the effectiveness of alternative deep feature extractors. For ground-level imagery, we additionally evaluate DINOv3 [114], BioCLIP [117] (a model trained to recognize over 450k diverse taxonomic labels), and embeddings extracted from MegaDetector [68]. We note that despite using blank camera trap images as input, these models might capture useful prior information based on the scene’s background. Supplemental Table B.2 shows macro-averaged normalized LPPD for those different models.

**Figure B.3:**
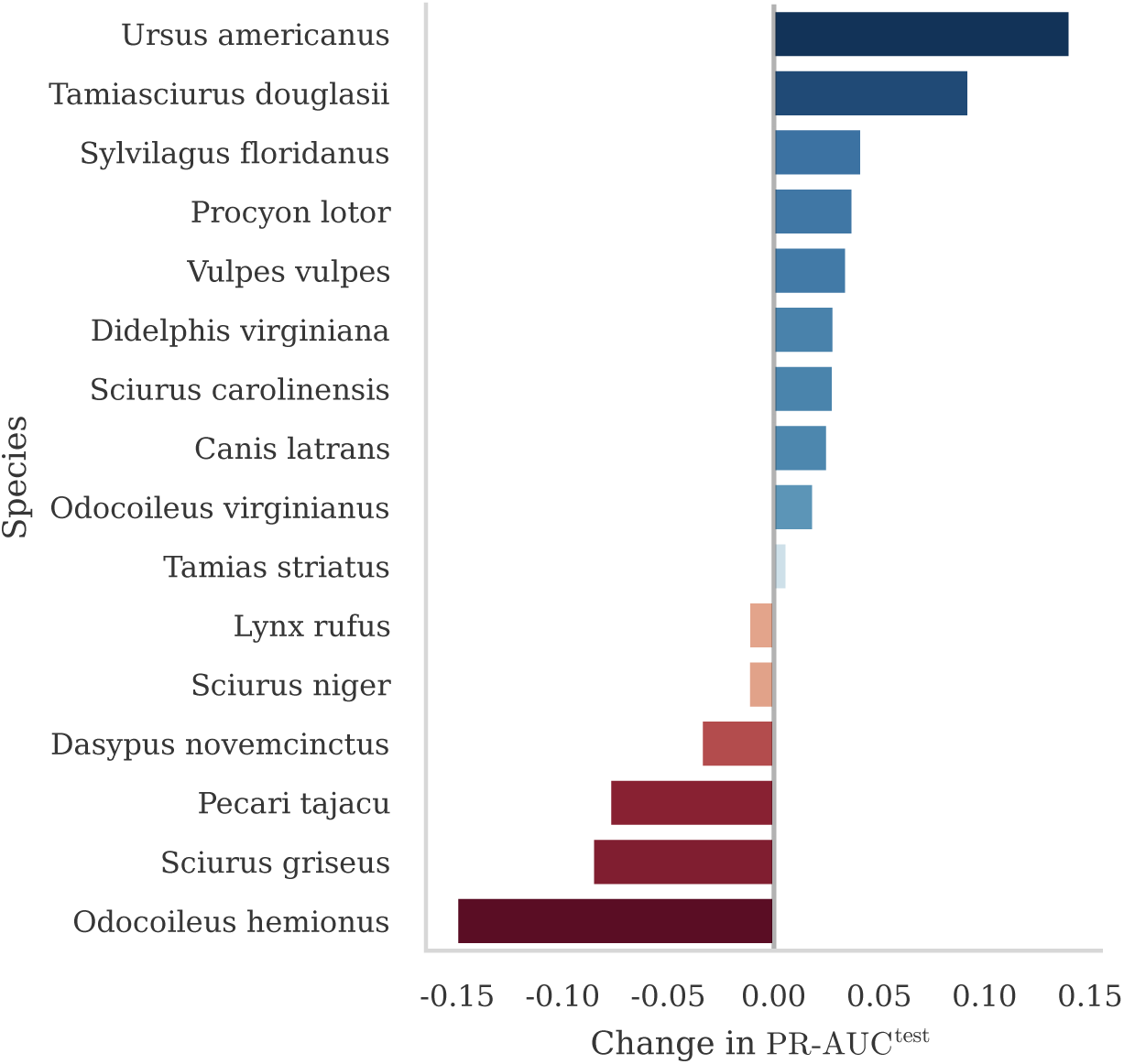
Test-set PR-AUC improvement of the best-performing model including satellite image features over the best-performing model not including satellite image features.

For satellite imagery, in addition to AlphaEarth Foundations [70], we evaluate Prithvi-EO-2.0 [118], SatBird [120], TaxaBind [113], Galileo [122], and TESSERA [81]. We furthermore evaluate DINOv3 pre-trained on satellite imagery [114], and the DINOv2 [104] general image model. We obtain embeddings from AlphaEarth Foundations [70] by sampling them from the Google Earth Engine Satellite Embedding V1 dataset. We use the annualized embedding from the year the sensor was deployed, at a spatial scale of 250 m, resulting in a single 64-dimensional feature vector per location. Supplemental Table B.3 shows predictive performance across those models.

**Table B.2:**
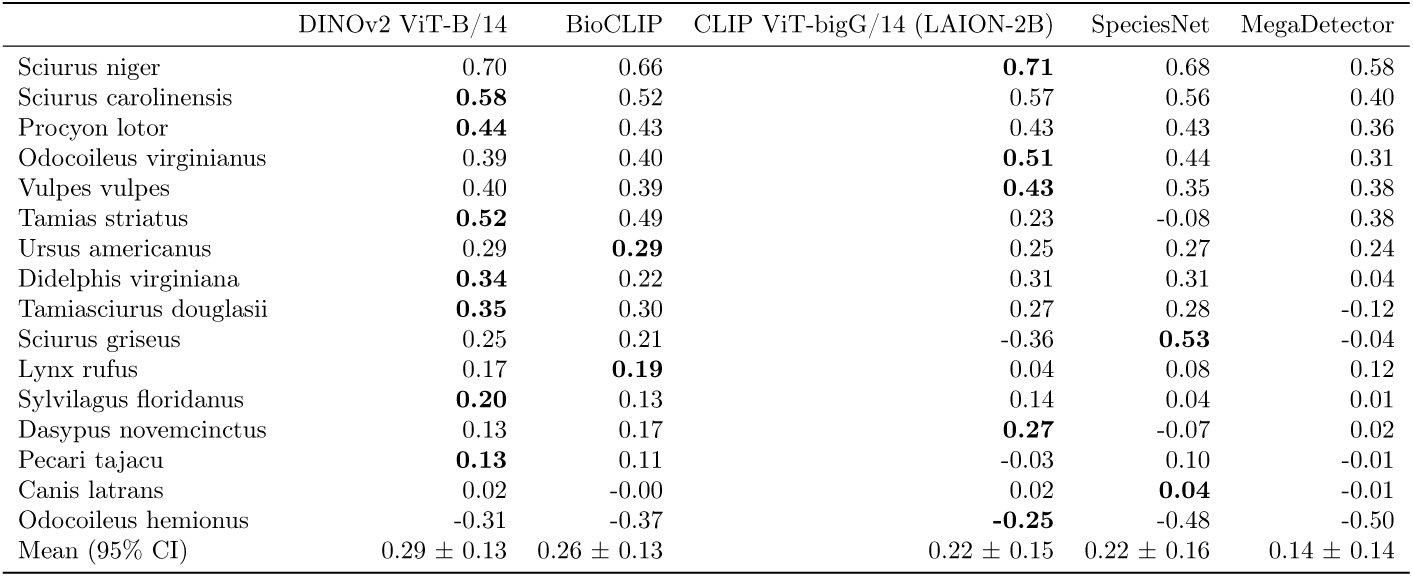
Normalized LPPD by ground-level image backbone and species. Best performance per species is bolded. **DINOv2 ViT-B/14 performs best overall, though the differences from the majority of other models are not significant.**

We note that while the features extracted by the majority of these models improve predictive performance by capturing *some* evidently meaningful habitat dimensions, they are not explicitly and solely optimized towards this task. This fact highlights significant potential for future work to adapt these models towards capturing more meaningful fine-grained habitat dimensions by adapting training objectives, training datasets, and potentially fine-tuning model weights explicitly for ecological modeling tasks.

### B.3 Habitat elements

As shown in Section 2.3, we can simplify occupancy models to use the CLIP-similarities between few natural-language habitat elements and ground-level images instead of using the full ground-level image features. Here we compare the predictive power of automatically extracted habitat elements (“VisDiff”, see Methods 4.4) and prior expert knowledge (“expert”, [71]). As illustrated by Supplemental Figure B.4, models using automatically-derived habitat elements perform similarly to the original full models for most but not all species. Some species even benefit slightly from simplified models, likely an artifact of decreased overfitting due to reduced model capacity. While expert-provided habitat elements seem to perform slightly worse, we note that one should only compare automatically-derived and expert-provided elements with caution. First, we obtained expert-provided habitat elements only for a subset of species. Second, the automatically-derived habitat elements are by definition optimized to be well-understood by the CLIP model, whereas this is not necessarily the case for the expert-provided elements. The full list of automatically-derived and expert-provided habitat elements can be found in Supplemental Table B.4.

**Table B.3:**
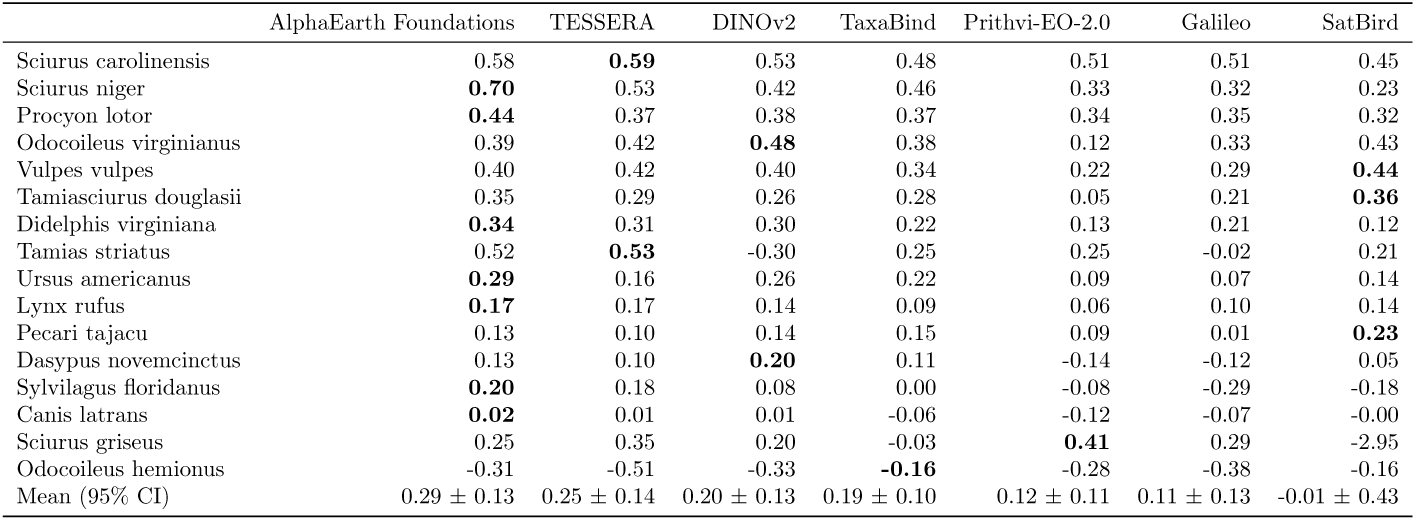
Normalized LPPD by satellite backbone and species. Best performance per species is bolded. AlphaEarth Foundations [70] performs best overall, though the differences to the majority of other models are not significant.

**Figure B.4:**
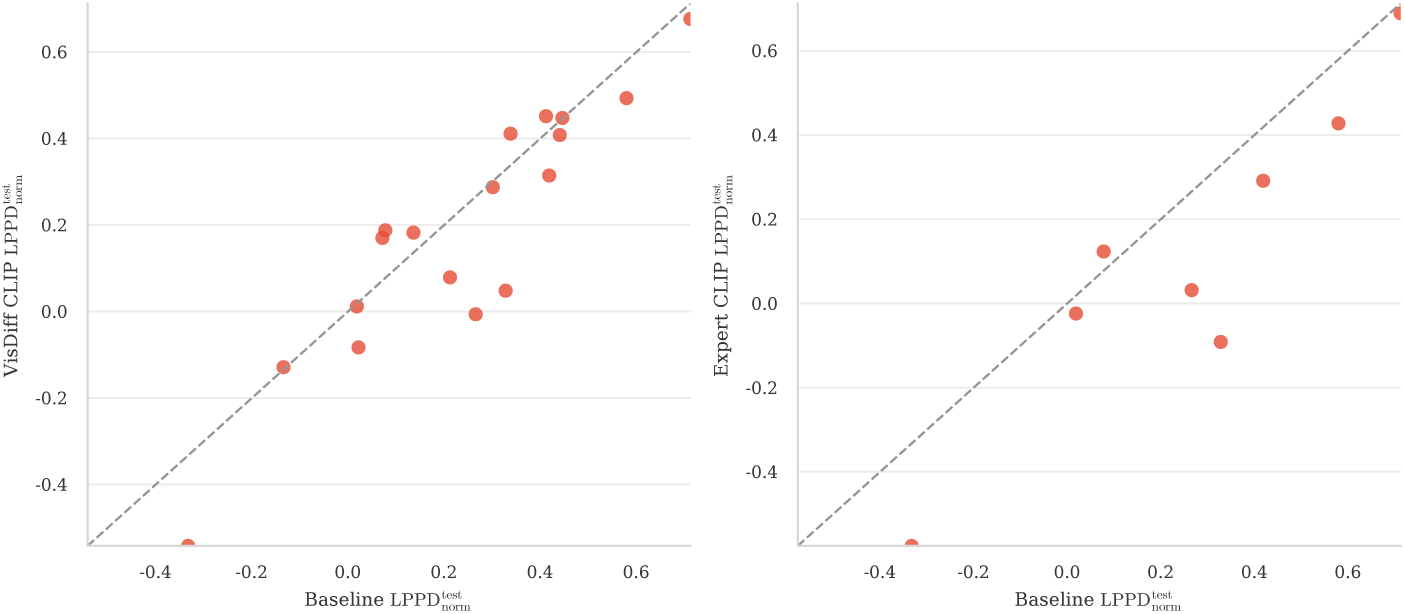
Predictive performance of models replacing the full ground-level image feature vector with automatically-derived (“VisDiff”, left) and expert-provided habitat elements (“expert”, right) vs. the baseline model which uses the entire feature vectors.

**Table B.4:**
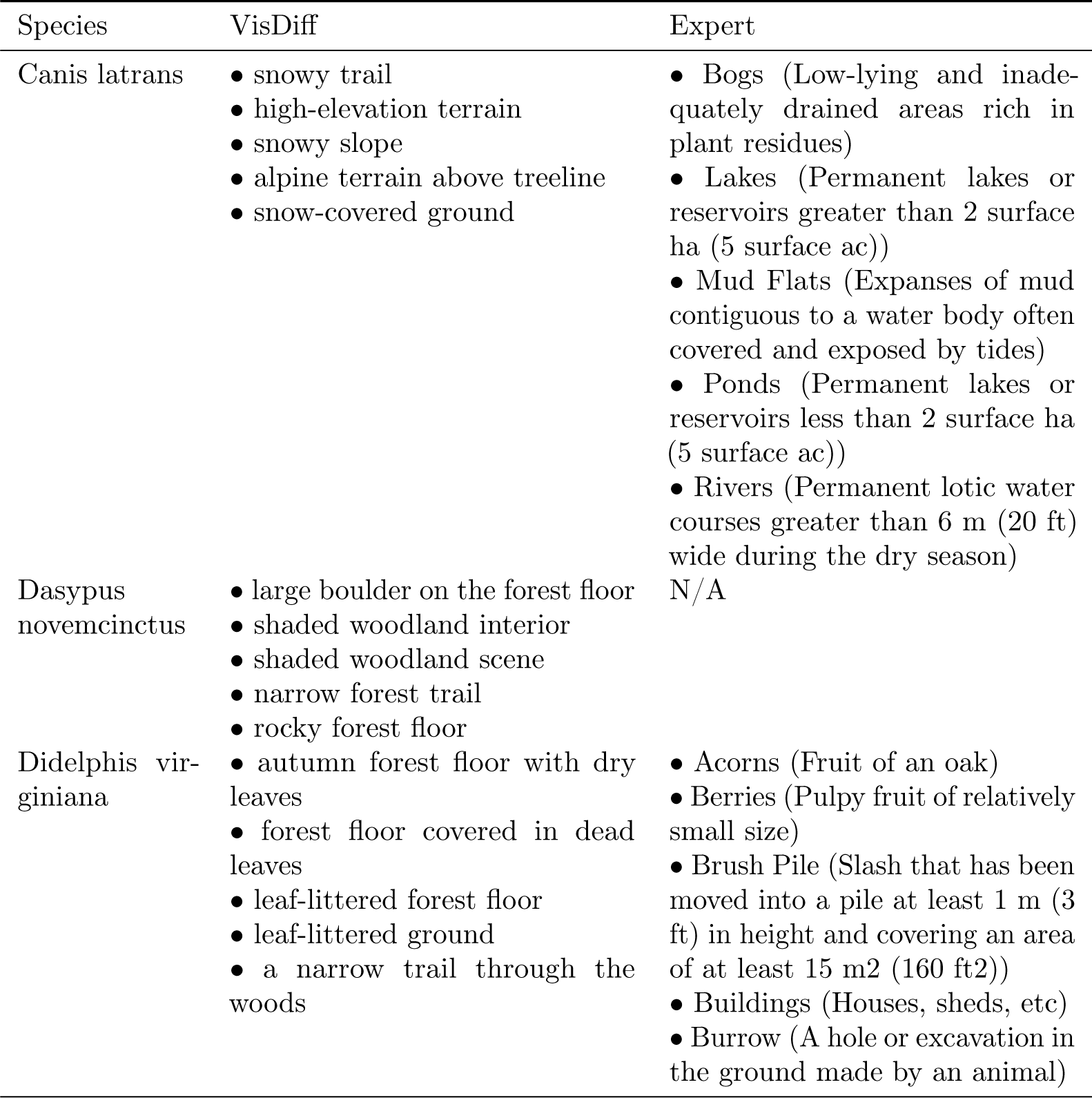
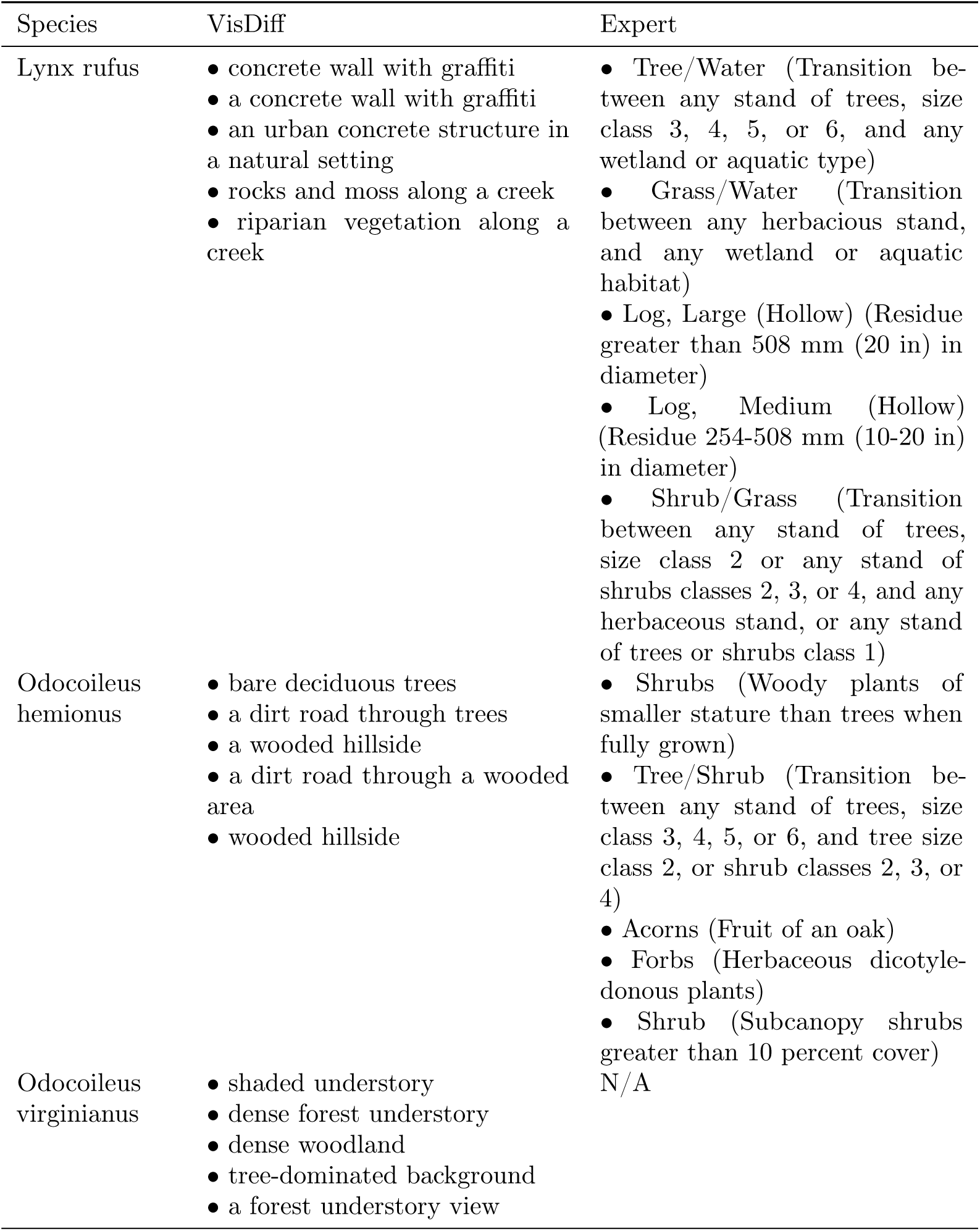
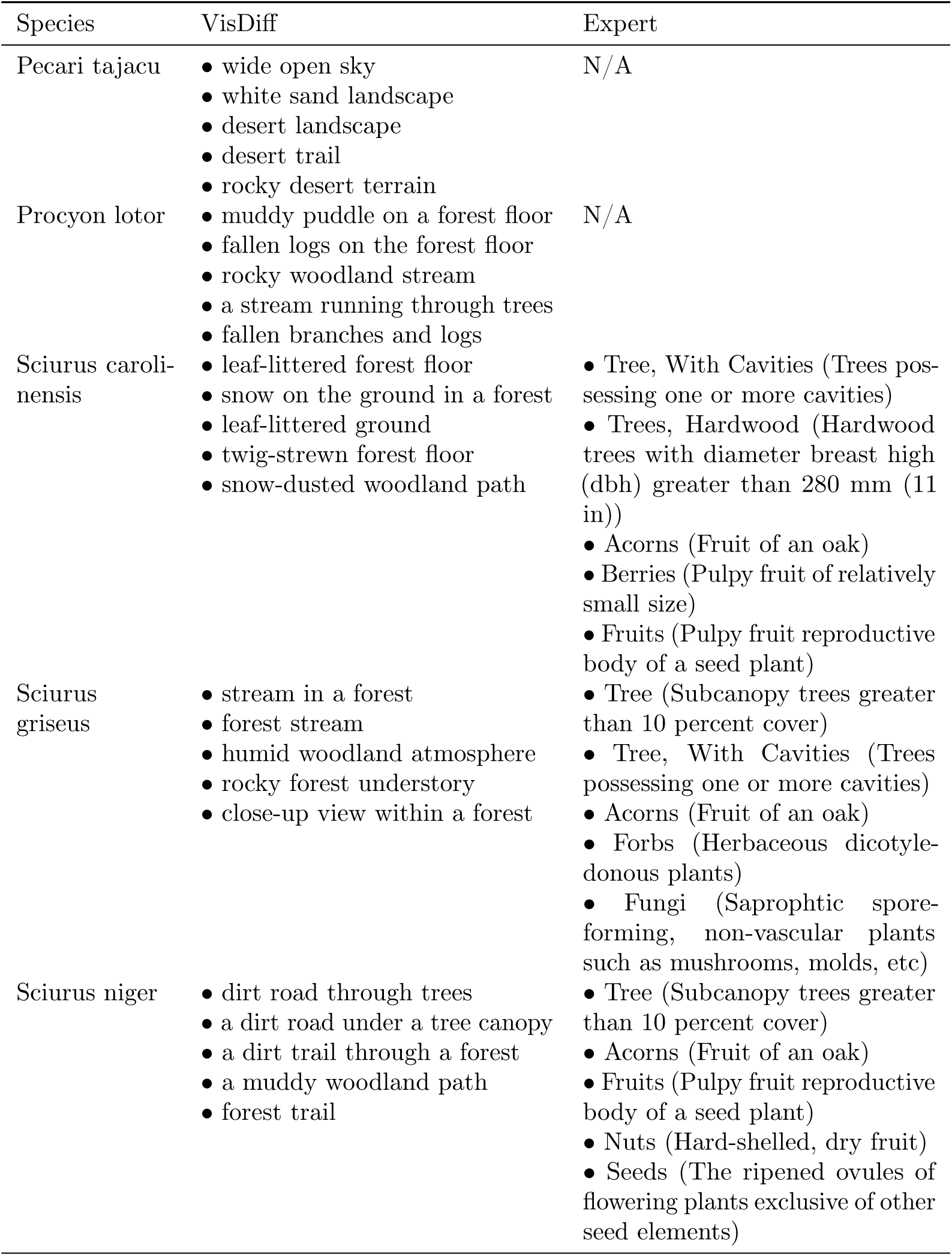
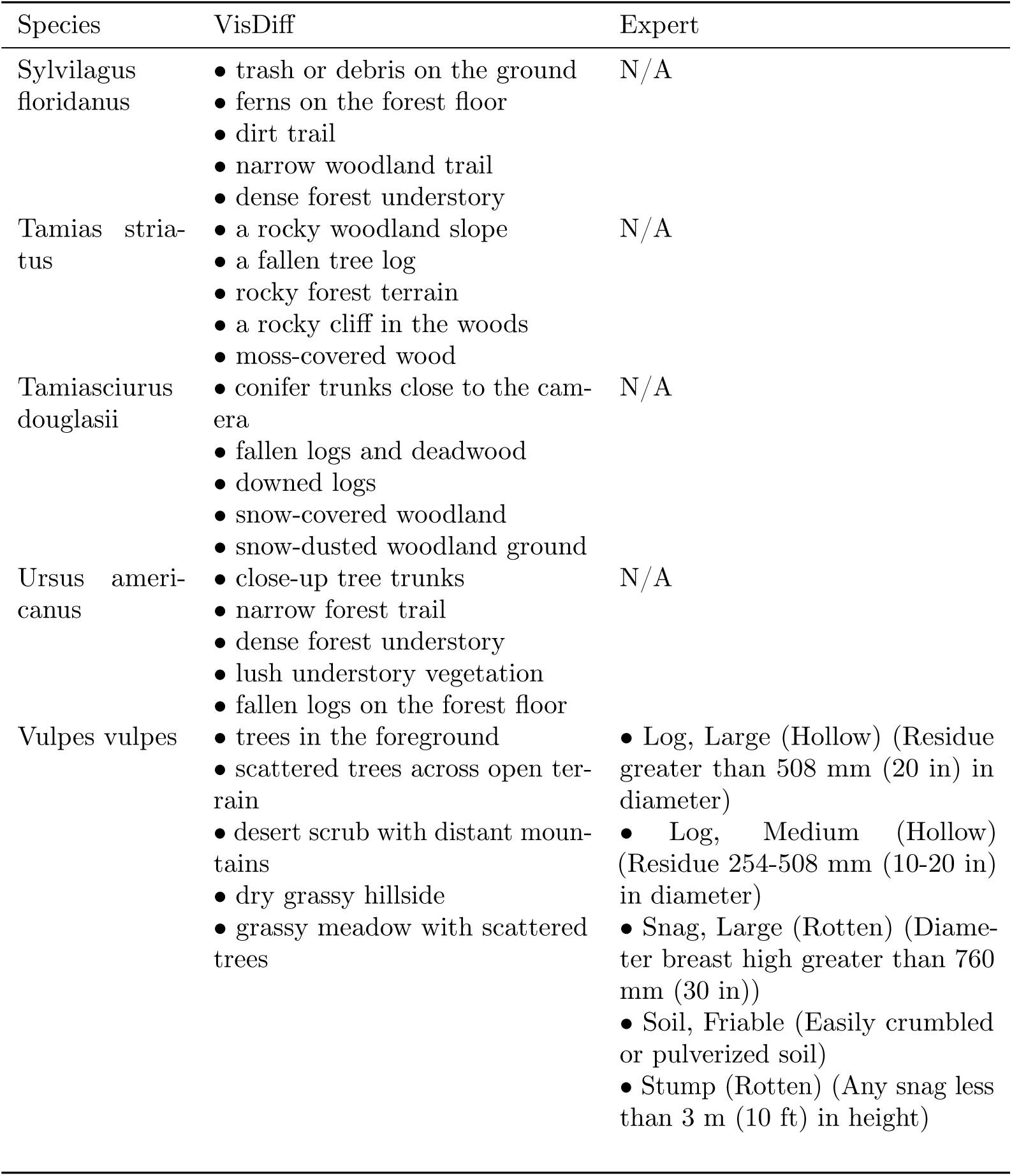
Habitat elements extracted automatically via VisDiff [80] and obtained from the California Wildlife Habitat Relationships System [71] (“Expert”). N/A signifies species missing from the aforementioned database.

| Species | VisDiff | Expert |
| --- | --- | --- |
| Canis latrans | • snowy trail<br>• high-elevation terrain<br>• snowy slope<br>• alpine terrain above treeline<br>• snow-covered ground | • Bogs (Low-lying and inadequately drained areas rich in plant residues)<br>• Lakes (Permanent lakes or reservoirs greater than 2 surface ha (5 surface ac))<br>• Mud Flats (Expanses of mud contiguous to a water body often covered and exposed by tides)<br>• Ponds (Permanent lakes or reservoirs less than 2 surface ha (5 surface ac))<br>• Rivers (Permanent lotic water courses greater than 6 m (20 ft) wide during the dry season) |
| Dasypus novemcinctus | • large boulder on the forest floor<br>• shaded woodland interior<br>• shaded woodland scene<br>• narrow forest trail<br>• rocky forest floor | N/A |
| Didelphis virginiana | • autumn forest floor with dry leaves<br>• forest floor covered in dead leaves<br>• leaf-littered forest floor<br>• leaf-littered ground<br>• a narrow trail through the woods | • Acorns (Fruit of an oak)<br>• Berries (Pulpy fruit of relatively small size)<br>• Brush Pile (Slash that has been moved into a pile at least 1 m (3 ft) in height and covering an area of at least 15 m<sup>2</sup> (160 ft<sup>2</sup>))<br>• Buildings (Houses, sheds, etc)<br>• Burrow (A hole or excavation in the ground made by an animal) |
| Continued on next page |  |  |
| Lynx rufus | • concrete wall with graffiti<br>• a concrete wall with graffiti<br>• an urban concrete structure in a natural setting<br>• rocks and moss along a creek<br>• riparian vegetation along a creek | • Tree/Water (Transition between any stand of trees, size class 3, 4, 5, or 6, and any wetland or aquatic type)<br>• Grass/Water (Transition between any herbaceous stand, and any wetland or aquatic habitat)<br>• Log, Large (Hollow) (Residue greater than 508 mm (20 in) in diameter)<br>• Log, Medium (Hollow) (Residue 254-508 mm (10-20 in) in diameter)<br>• Shrub/Grass (Transition between any stand of trees, size class 2 or any stand of shrubs classes 2, 3, or 4, and any herbaceous stand, or any stand of trees or shrubs class 1) |
| Odocoileus hemionus | • bare deciduous trees<br>• a dirt road through trees<br>• a wooded hillside<br>• a dirt road through a wooded area<br>• wooded hillside | • Shrubs (Woody plants of smaller stature than trees when fully grown)<br>• Tree/Shrub (Transition between any stand of trees, size class 3, 4, 5, or 6, and tree size class 2, or shrub classes 2, 3, or 4)<br>• Acorns (Fruit of an oak)<br>• Forbs (Herbaceous dicotyledonous plants)<br>• Shrub (Subcanopy shrubs greater than 10 percent cover) |
| Odocoileus virginianus | • shaded understory<br>• dense forest understory<br>• dense woodland<br>• tree-dominated background<br>• a forest understory view | N/A |
| Pecari tajacu | • wide open sky<br>• white sand landscape<br>• desert landscape<br>• desert trail<br>• rocky desert terrain | N/A |
| Procyon lotor | • muddy puddle on a forest floor<br>• fallen logs on the forest floor<br>• rocky woodland stream<br>• a stream running through trees<br>• fallen branches and logs | N/A |
| Sciurus carolinensis | • leaf-littered forest floor<br>• snow on the ground in a forest<br>• leaf-littered ground<br>• twig-strewn forest floor<br>• snow-dusted woodland path | • Tree, With Cavities (Trees possessing one or more cavities)<br>• Trees, Hardwood (Hardwood trees with diameter breast high (dbh) greater than 280 mm (11 in))<br>• Acorns (Fruit of an oak)<br>• Berries (Pulpy fruit of relatively small size)<br>• Fruits (Pulpy fruit reproductive body of a seed plant) |
| Sciurus griseus | • stream in a forest<br>• forest stream<br>• humid woodland atmosphere<br>• rocky forest understory<br>• close-up view within a forest | • Tree (Subcanopy trees greater than 10 percent cover)<br>• Tree, With Cavities (Trees possessing one or more cavities)<br>• Acorns (Fruit of an oak)<br>• Forbs (Herbaceous dicotyledonous plants)<br>• Fungi (Saprophytic spore-forming, non-vascular plants such as mushrooms, molds, etc) |
| Sciurus niger | • dirt road through trees<br>• a dirt road under a tree canopy<br>• a dirt trail through a forest<br>• a muddy woodland path<br>• forest trail | • Tree (Subcanopy trees greater than 10 percent cover)<br>• Acorns (Fruit of an oak)<br>• Fruits (Pulpy fruit reproductive body of a seed plant)<br>• Nuts (Hard-shelled, dry fruit)<br>• Seeds (The ripened ovules of flowering plants exclusive of other seed elements) |
| Continued on next page |  |  |
| Sylvilagus floridanus | • trash or debris on the ground<br>• ferns on the forest floor<br>• dirt trail<br>• narrow woodland trail<br>• dense forest understory | N/A |
| Tamias striatus | • a rocky woodland slope<br>• a fallen tree log<br>• rocky forest terrain<br>• a rocky cliff in the woods<br>• moss-covered wood | N/A |
| Tamiasciurus douglasii | • conifer trunks close to the camera<br>• fallen logs and deadwood<br>• downed logs<br>• snow-covered woodland<br>• snow-dusted woodland ground | N/A |
| Ursus americanus | • close-up tree trunks<br>• narrow forest trail<br>• dense forest understory<br>• lush understory vegetation<br>• fallen logs on the forest floor | N/A |
| Vulpes vulpes | • trees in the foreground<br>• scattered trees across open terrain<br>• desert scrub with distant mountains<br>• dry grassy hillside<br>• grassy meadow with scattered trees | • Log, Large (Hollow) (Residue greater than 508 mm (20 in) in diameter)<br>• Log, Medium (Hollow) (Residue 254-508 mm (10-20 in) in diameter)<br>• Snag, Large (Rotten) (Diameter breast high greater than 760 mm (30 in))<br>• Soil, Friable (Easily crumbled or pulverized soil)<br>• Stump (Rotten) (Any snag less than 3 m (10 ft) in height) |

## C Experimental details

### C.1 Environmental variables

Covariate selection in ecological models is notoriously difficult and oftentimes driven by prior species-specific expertise about which environmental variables are ecologically relevant. Due to the large number of species considered in this work, and in order to make results comparable across species, we relied on a general set of environmental variables that are widespread and well-proven in the species distribution modeling literature [120]. Supplemental Table C.1 shows the full set of environmental variables we use as occupancy covariates.

**Table C.1:**
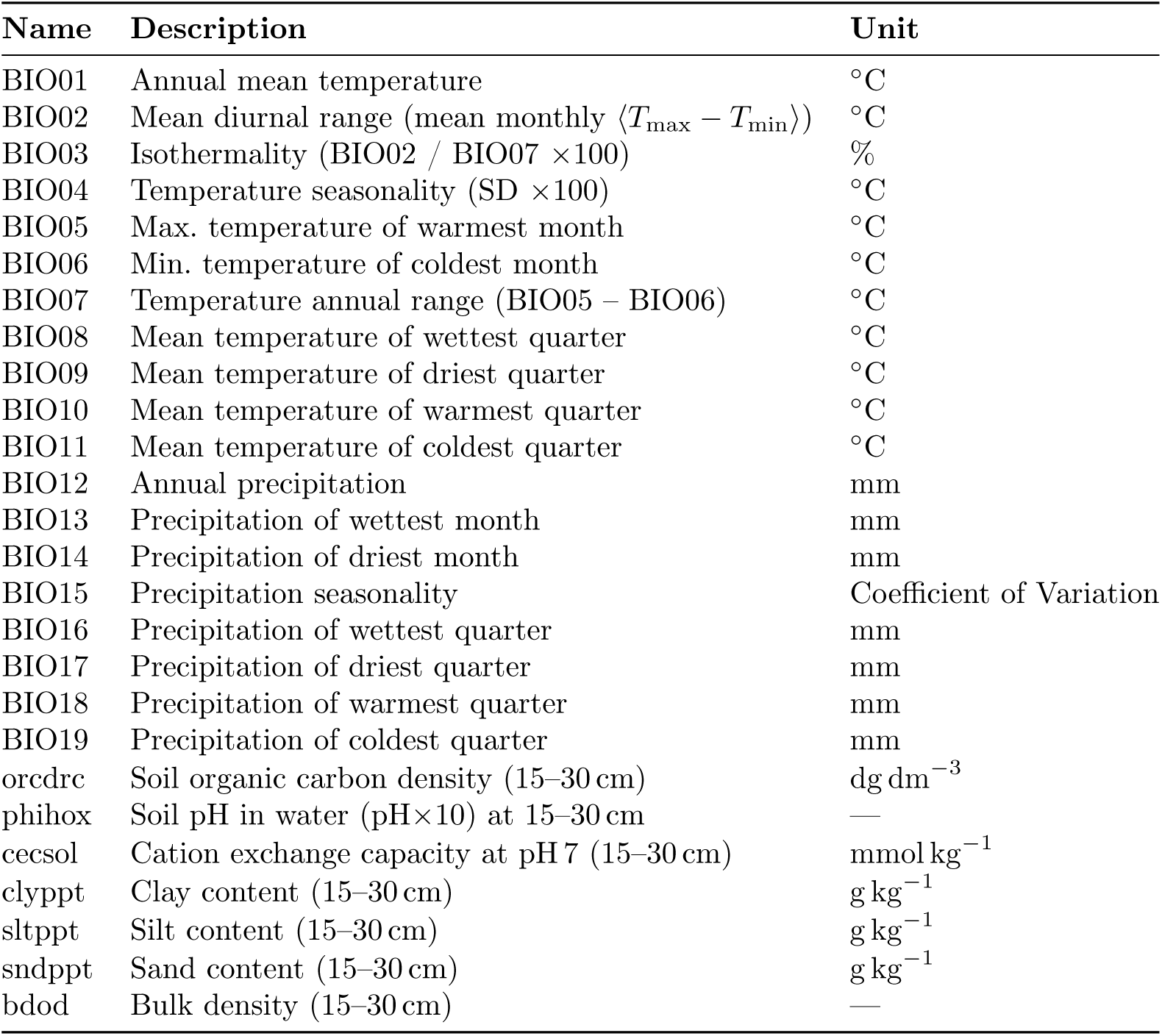
The full set of environmental variables used for our experiments, consisting of WorldClim BIO [90] and SoilGrids [108] variables sampled via Google Earth Engine. We adapted this set of environmental variables from SatBird [120] but excluded the SoilGrids “absolute depth to bedrock in cm” since it is not readily available on Google Earth Engine.

### C.2 Technical implementation

To support the experiments presented in this manuscript, we developed *Biolith*, a new implementation of occupancy and related statistical ecological models in the Python programming language. Since Python is the lingua franca of machine learning research, we argue that this reduces the barrier of applying new machine learning algorithms to ecological modeling problems. Biolith is highly influenced by popular libraries in the statistical ecology ecosystem, including Unmarked [82, 97] and spOccupancy [78] in particular.

Beyond the classic Bernoulli occupancy model by [101] used in this manuscript, Biolith implements the abundance model by [111], the continuous-score occupancy model by [109] for incorporating uncertain machine learning classifications, and the discrete-time counting occurrences model by [105]. In contrast to existing implementations, Biolith places great emphasis on modularity; all of the models named above can optionally be extended with modules such as false positives (thereby accounting for faulty observations of *presence*, which is useful when incorporating classifications produced by machine learning models) [110], site-and observation-level random effects, and spatial explicitness to model spatial autocorrelation.

Biolith is implemented using the NumPyro [106] probabilistic programming library, which enables defining models declaratively in a fashion similar to languages such as BUGS / JAGS. To fit models, Biolith uses the NumPyro implementation of Hamiltonian Monte Carlo inference and the No-U-Turn Sampler [91].

Since NumPyro is implemented using JAX [69], inference can be accelerated by graphics processing units (GPUs). The massive parallelism of GPUs unlocks potential reductions in inference runtime; however, currently this manifests itself exclusively on large datasets (see Supplemental Figure C.1). We perform all of our experiments on 16 cores of an AMD EPYC 9554 with 64 GB of system memory and a single NVIDIA L40S GPU with 48 GB of memory. We use JAX 0.5.2, NumPyro 0.18.0, Biolith 0.0.9, and CUDA 12.9.

**Figure C.1:**
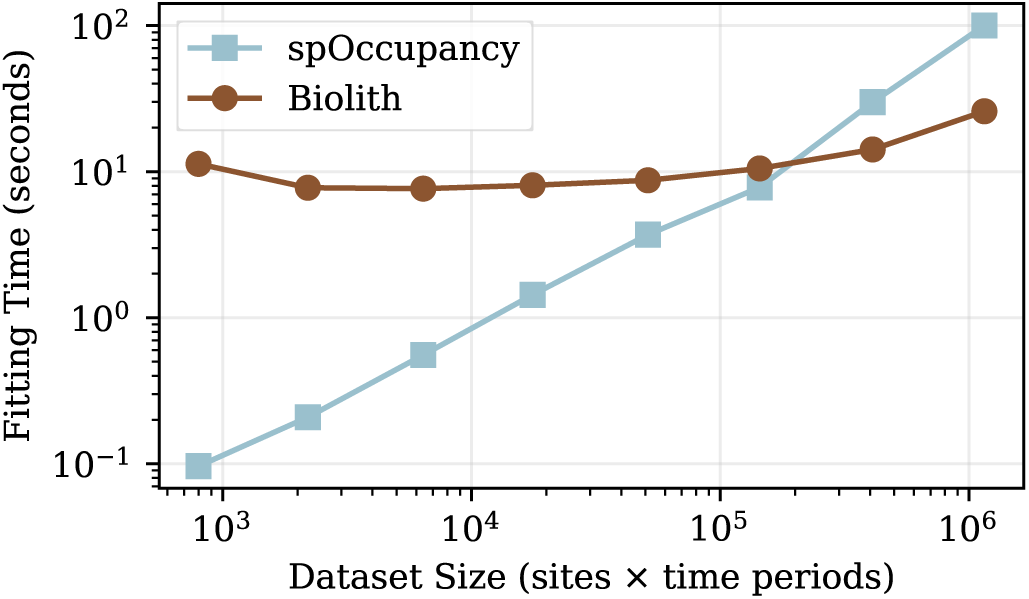
Benchmark of Biolith (GPU-accelerated) and spOccupancy [78] (CPU-only, single-threaded) runtimes over different dataset sizes. spOccupancy shows an almost perfect linear relationship and performs best at small to moderate dataset sizes, whereas the runtime of Biolith is initially strongly dominated by constant overhead and only becomes competitive on larger datasets. Details about the system used to run this benchmark can be found in Section C.2.

